# Targeting neuronal and glial cell types with synthetic promoter AAVs in mice, non-human primates, and humans

**DOI:** 10.1101/434720

**Authors:** Josephine Jüttner, Arnold Szabo, Brigitte Gross-Scherf, Rei K. Morikawa, Santiago B. Rompani, Miguel Teixeira, Peter Hantz, Tamas Szikra, Federico Esposti, Cameron S. Cowan, Arjun Bharioke, Claudia P. Patino-Alvarez, Özkan Keles, Chiara N. Roth, Akos Kusnyerik, Nadin Gerber-Hollbach, Thierry Azoulay, Dominik Hartl, Arnaud Krebs, Dirk Schübeler, Rozina Hajdu, Akos Lukats, Janos Nemeth, Zoltan Z. Nagy, Kun-Chao Wu, Rong-Han Wu, Lue Xiang, Xiao-Long Fang, Zi-Bing Jin, David Goldblum, Pascal W. Hasler, Hendrik Scholl, Jacek Krol, Botond Roska

## Abstract

Targeting genes to specific neuronal or glial cell types is valuable both for understanding and for repairing brain circuits. Adeno-associated viral vectors (AAVs) are frequently used for gene delivery, but targeting expression to specific cell types is a challenge. We created a library of 230 AAVs, each with a different synthetic promoter designed using four independent strategies. We show that ~11% of these AAVs specifically target expression to neuronal and glial cell types in the mouse retina, mouse brain, non-human primate retina *in vivo*, and in the human retina *in vitro*. We demonstrate applications for recording, stimulation, and molecular characterization, as well as the intersectional and combinatorial labeling of cell types. These resources and approaches allow economic, fast, and efficient cell-type targeting in a variety of species, both for fundamental science and for gene therapy.

## Introduction

The concept of cell types – morphologically, physiologically, and molecularly similar groups of neurons or glia within a given brain region – has become an important starting point for understanding and modulating brain function. In basic research, genetic labeling of neuronal or glial cell types enables their isolation and molecular characterization. Genetically encoded sensors or electrical recording targeted to neuronal cell types allow monitoring of activity, and cell-type-targeted optogenetic or chemogenetic tools enable modulation of their activity. In translational research, cell-type-targeted modulation of brain function and cell-type-specific gene replacement are repair strategies for treating human diseases.

Despite the central importance for both basic and translational research, most current technologies available for cell-type-targeting rely on transgenic animals, which limits their applicability. Either the genetic tool that senses or modulates brain function, or the enzyme, such as Cre recombinase, that allows the genetic tool to be conditionally expressed, is expressed from the animal’s genome. The inclusion of a transgenic component in the cell-type-targeting strategy excludes its use in therapy for humans, limits its range of application in pre-clinical, non-human primate research, and complicates its use in model organisms such as mice. The development of transgenic non-human primates and mice is costly and slow, especially since cell-type targeting is often applied in the context of other genetic manipulations, such as double or triple gene knockouts, or when targeting different cell types with different tools.

Viral vectors for cell-type-targeting may overcome such limitations. AAVs are the most frequently used vectors in both basic research and gene therapy, as they are safe for use in all tested species, including humans and non-human primates, and their production is simple, cheap, and fast (Planul and Dalkara, 2017). They have three important components: the capsid for cell entry, the promoter that drives transgene expression, and the gene of interest to be expressed in the transduced cells, and they drive expression episomally (Duan et al., 1998; Penaud-Budloo et al., 2008). Futhermore, many genetic tools are small enough to fit into AAVs, different AAVs can be injected together, and synthetic AAV capsids allow brain-wide delivery (Deverman et al., 2016).

Cell-type-targeting by AAVs could be achieved by engineering the capsid and/or by using specific promoters. Capsid protein mutations can be used to tune the efficacy of AAVs for entry into a variety of cell types, but they rarely result in cell-type-specific entry (Dalkara et al., 2013; Cronin et al., 2014; Chan et al., 2017). AAVs equipped with cell-type-specific promoters are potentially useful for targeting expression to cell types; however, existing promoters in AAVs, with a few exceptions (Oh et al., 2009; Khabou et al., 2018), target expression only broadly in a collection of cell types, such as inhibitory neurons in the brain, or retinal ganglion or ON bipolar cells (Cronin et al., 2014; Nathanson et al., 2009; Hanlon et al., 2017; Dimidschstein et al., 2016). It is not clear whether promoter-dependent targeting can be generalized to many individual cell types. One way to search for AAVs targeting specific cell types is to screen a library of promoters driving transgene expression in brain regions of interest. An AAV promoter screen for targeting cell types has not yet been performed.

We have developed a library of 230 AAVs, each with a different synthetic promoter, most of them (226) driving an optogenetic tool fused to a fluorescent marker. The marker-tagged optogenetic tool enables cell types to be both identified and manipulated. First we tested the AAVs for cell-type-specific expression in the eyes of mice *in vivo*. We then analyzed a subset in the brains of mice *in vivo* (*n* = 48 AAVs), in the eye of primates *in vivo* (*n* = 94), and in human post-mortem retinas *in vitro* (*n* = 84). Remarkably, ~11% of the AAVs tested drove expression in individual cell types, and many others in combinations of cell types. The cell types targeted by a specific synthetic promoter varied widely across mouse and primate retinas, but less across non-human primate and human retinas. We created logic OR and AND gates using combinations of AAVs to target cell types that could not be marked by any of the AAVs alone. Finally, we show the use of cell-type-specific AAVs for molecular analysis and for manipulating and recording neuronal function.

Our results demonstrate that different neuronal and glial cell types of mice, non-human primates, and humans can be efficiently targeted using AAVs. Furthermore, we describe a set of AAVs applicable in basic research for recording or modulating the activities of cell types, and in translational research for gene therapy of cell-type-specific human diseases such as retinitis pigmentosa and macular degeneration.

## Results

### AAV-based cell-type-targeting strategy

We created a library of 230 AAV plasmids, each equipped with a different synthetic DNA sequence of mean length 1249±673 bp in the range of 113-2501 nt (Figure S1A and Tables S1-S3), positioned 5’ of a transgene. The 5’ sequences (“synthetic promoters”) were constructed using four different strategies (A-D). The synthetic promoter group ProA included sequences upstream of the start codon of selected mouse retinal cell-type-specific genes (Siegert et al., 2012). Group ProB was generated by ordered an assembly of phylogenetically conserved DNA elements identified in a nucleotide sequence preceding the transcription initiation sites of a minimum of two genes with the highest cell specificity and expression indices (Siegert et al., 2012). ProC was made up of repeated transcription factor binding sites of cell-type-specific transcription factors (Siegert et al., 2012) interleaved with random sequences. ProD was identified based on an approach combining epigenome and transcriptome profiling, and consisted of low-methylated *cis*-regulatory elements transcriptionally active in different retinal cell types (Hartl et al., 2017). ProC and ProD also contained a minimal TATA-box promoter element (Figure 1A).

**Figure 1.**
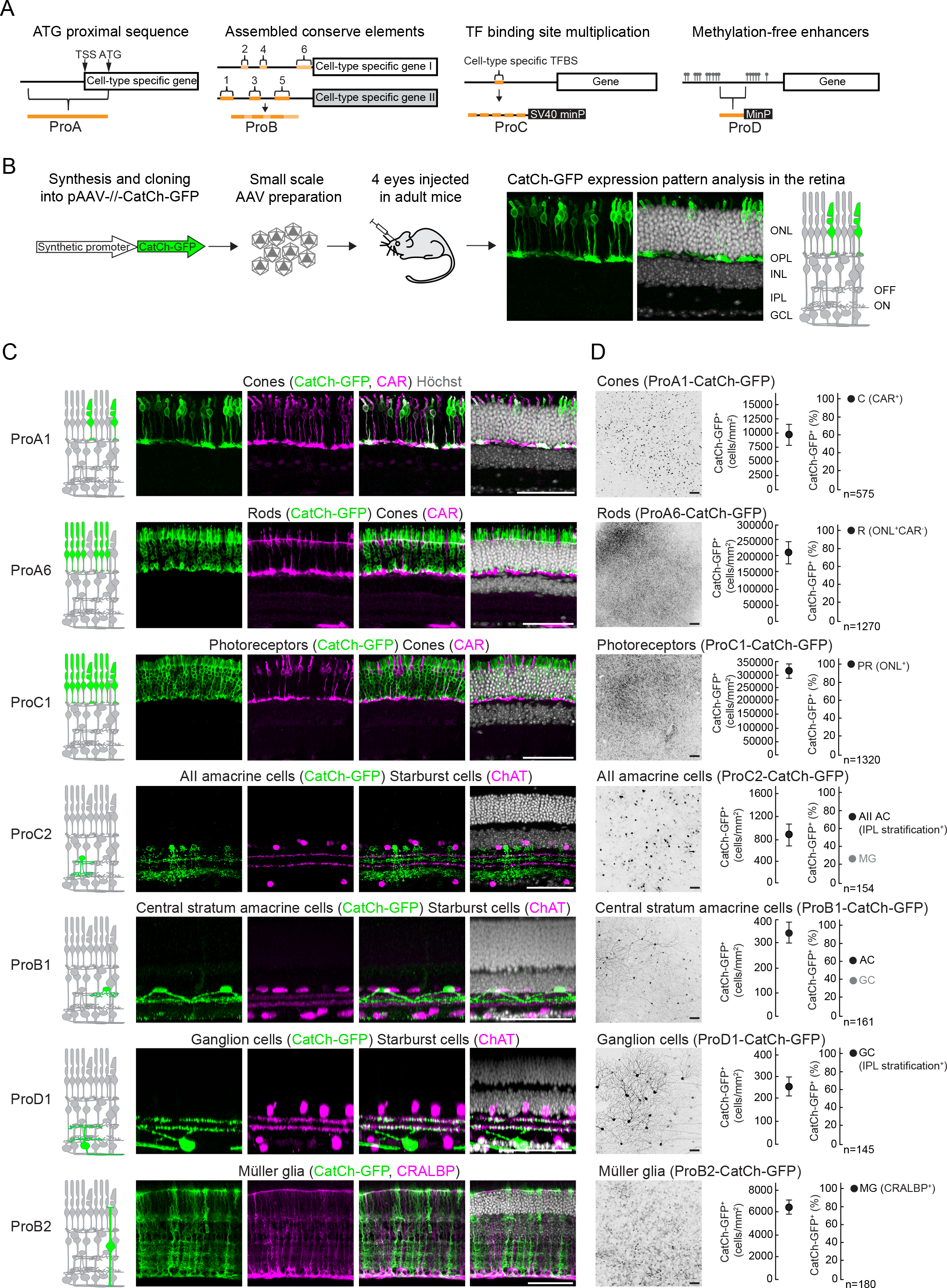
Cell-type targeting in mouse retina *in vivo.* (A) Synthetic promoter design strategies. TSS, transcription start site; ATG, translation start site; TF, transcription factor; TFBS, transcription factor binding site; minP, minimal promoter. (B) Workflow for AAV-based retinal cell-type targeting screen. (C) Confocal images of sections of AAV-infected retinas. Left, CatCh-GFP (green); middle-left, immunostaining with marker (magenta) indicated above; middle-right, CatCh-GFP and marker; right, CatCh-GFP and marker and nuclear stain (Höchst, white). Scale bars, 50 μm. (D) Left, confocal images of AAV-infected retinas (top view), CatCh-GFP (black). Middle, quantification of CatCh-GFP^+^ cell density, values are means ± SEM from images taken from four retinas. Right, quantification of AAV-targeting specificity shown as a percentage of the major (black) and minor (grey) cell types or classes among cells expressing CatCh-GFP. C, cones; R, rods; PR, photoreceptors; AII AC, AII amacrine cells; MG, Müller glia; AC, amacrine cells; GC, ganglion cells. Scale bars, 50 μm. See also Figure S1 and Table S1.

Of the 230 AAVs, 226 were designed to drive a channelrhodopsin variant fused to GFP (CatCh-GFP) (Kleinlogel et al., 2011). The use CatCh-GFP in the screen was based on two rationales. First, we found that expression of GFP alone was much higher than the optogenetic tool when driven by the same synthetic promoter. Since the use of the synthetic promoter might involve optogenetics, we selected fewer but powerful synthetic promoters that could drive hard-to-express genes. Second, the cell membrane-bound CatCh-GFP protein revealed the fine morphology of neuronal processes better than diffusible markers such as GFP. The remaining four AAVs were designed to express GFP.

AAV capsids are preferentially infectious for different host cells. We used four different AAV capsids to adapt to different species and infection routes: AAV8 (Allocca et al., 2007), AAV9 (Allocca et al., 2007; Lebherz et al., 2008), and AAV8BP2 (Cronin et al., 2014) for adult mouse retina; AAV8BP2 for adult non-human primate, and human retina; AAV9 (Aschauer et al., 2013) for adult mouse cortex; and AAV-PHP.B (Deverman et al., 2016) for brain-wide labeling via intravenous administration (Table S1-S3).

We developed a “rapid-AAV” protocol that increased the speed of production 10-fold. “Rapid-AAVs” were used for the *in vivo* screens in mouse retina and cortex (Figure 1B). The screens in non-human primates and humans required substantial amounts of viral vector, and were performed with AAVs produced conventionally (Grieger et al., 2006a).

### Cell-type targeting in mouse retina *in vivo*

We injected 920 mouse eyes subretinally with the 230 AAVs, four eyes with each AAV, and evaluated transgene expression 3-4 weeks later using confocal microscopy of whole-mount retinas (Figure 1B). Of the 230 synthetic promoters tested, 113 induced transgene expression in at least one retinal cell type, with ProA, ProB, and ProC promoters showing similar target efficiency (Figure S1B and Table S1). Twenty five synthetic promoters led to expression in specific cell types and many others labeled a combination of selected cell types (Table S1). Despite this specificity, less than 1% of synthetic promoters replicated the expression specificity of their source genes. Two synthetic promoters, ProA1 and ProA4, drove CatCh-GFP expression specifically in cone photoreceptors, identified by the characteristic position of the cell bodies at the outer margin of the retinal outer nuclear layer (ONL) and co-labeling with the cone-specific marker cone arrestin (CAR) (Zhu et al., 2002) (Figures 1C and S1C). Nine synthetic promoters, ProA6, ProB5, ProC22, ProC32, ProD2, ProD3, ProD4, ProD5, and ProD6, were found to be rod specific, driving CatCh-GFP expression in CAR-negative ONL cells (Figures 1C and S1C and Table S1). ProC1 targeted transgene expression to both photoreceptor types (Figure 1C).

For inner retinal neurons, we identified several synthetic promoters that labeled amacrine and ganglion cell types. These cells can be distinguished by their overall morphology and the stratification of their processes at different depths in the retinal inner plexiform layer (IPL) (Masland, 2001; Wässle, 2004). A subset of amacrine cells (starburst cells) with processes in two thin strata in the IPL were labeled with an antibody against choline acetyltransferase (ChAT) to provide a depth marker (Haverkamp and Wässle, 2000). Two synthetic promoters, including ProC2, induced transgene expression in AII amacrine cells and a few Müller glia cells (MacNeil and Masland, 1998) (Figure 1C). Synthetic promoters such as ProB1 targeted other types of amacrine cells with processes located in one stratum of the IPL (Figure 1C). Several synthetic promoters were identified that targeted different ganglion cell types with distinguishable stratification; for example, ProD1 targeted a set of bistratified ganglion cells with dendrites tightly aligned with those of ChAT-positive amacrine cells (Figure 1C and Table S1). Selective targeting of bipolar cells was rare; among the AAVs tested only AAV-ProB4 resulted in transgene expression in a type of OFF bipolar cells and cones (Table S1). Our screen also yielded AAVs targeting non-neuronal cells. The synthetic promoters ProB2, ProA3, ProA18, ProA21, ProA22, ProC17, or ProD17 induced CatCh-GFP expression in Müller glial cells, co-labeled with the specific marker CRALBP (Sarthy et al., 1998) (Figure 1C and Table S1).

To define the cell-type-targeting efficiency of selected synthetic promoters, we quantified the density of CatCh-GFP^+^ cells relative to the overall density of cells of a particular cell type. ProA1, the most effective cone-targeting synthetic promoter, highlighted on average 9,950±2,110 of 12,000 CAR^+^ cells per mm^2^, i.e., ~83% of adult cones (Jeon et al., 1998; Ortín-Martínez et al., 2014) (Figure 1D). The targeting efficiency of AAV-ProA6 was ~50% of rods (~ 217,600±32,450 CatCh-GFP^+^ cells out of 437,000 cells per mm^2^) (Jeon et al., 1998), whereas photoreceptor-targeting AAV-ProC1 highlighted 68% of the ONL cells. AAV-ProC2 targeted ~34% of AII amacrine cells expressing the specific marker Dab1 (Rice and Curran, 2000), whereas ProD1 produced transgene expression in 37% of ganglion cells (250±48 out of 672± 23 cells per mm^2^) that had dendrites within both ChAT strata (Sun et al., 2002; Salinas-Navarro et al., 2009). AAV-ProB2 highlighted ~44% of Müller glia expressing CRALBP (Figure 1D).

We next quantified target specificity by determining the percentage of one cell type or cell class in the overall CatCh-GFP^+^ cell population highlighted by a particular AAV. Remarkably, based on morphology or marker expression, several AAVs (for example AAV-ProA1, -ProA6, -ProD1, -ProB2, -ProA4, and -ProD2) were almost fully cell-type specific (Figures 1D and S1D and Table S1). For other AAVs with highly restricted expression patterns, particularly those targeting inner-retinal neurons, the target specificity was reduced by co-expression in another cell type, often Müller glia (for example AAV-ProC2, -ProC6, -ProA3). Co-labeling could potentially be eliminated using an intersectional strategy (see below). In other cases, members of a single cell class were labeled, such as the photoreceptor cell class by AAV-ProC1 or the ganglion cell class by ProA5 (Figure S2).

Some applications require sparse targeting of neurons of a given type, and several AAVs targeted particular retinal cell types sparsely (Table S1). For example, AAV-ProD2 produced expression in 10,025±1,250 rods per mm^2^, AAV-ProC6 targeted 250±57 AII amacrine cells per mm^2^, while AAV-ProA3 infected 463±156 ganglion cells per mm^2^ stratified in the IPL OFF-sublaminae (Figure S1C,D).

Taken together, screening in mouse retina identified a variety of synthetic promoters introduced into AAVs that targeted transgenes to specific mouse retinal cell types either efficiently or sparsely.

### AND/OR logic for cell-type targeting

Besides AAVs targeting individual cell types, our screen also identified AAVs that targeted two or more distinct retinal cell types simultaneously (Table S1). The overlap between cell types targeted by different AAVs provided an intersectional strategy (logic AND gate) to target a cell type, such as horizontal cells, for which no specific synthetic promoter was identified (Figure 2A). To express CatCh-GFP in horizontal cells, we leveraged two AAVs with different synthetic promoters, ProB3 and ProC3, targeting photoreceptors/horizontal cells and horizontal/amacrine/ganglion cells, respectively (Figure 2B). For intersectional transgene expression, AAV-ProB3 carried a Cre-dependent double-inverted (DIO) CatCh-GFP coding sequence and AAV-ProC3 drove expression of Cre recombinase and a fluorescent mCherry marker. CatCh-GFP-expression was not induced by infection of retina with AAV-ProB3-DIO-CatCh-GFP alone. Infection solely with AAV-ProC3-Cre-mCherry produced a mixture of mCherry-labeled cell types according to ProC3 specificity. In retinas co-injected with both AAVs, CatCh-GFP was expressed only in horizontal cells, with concomitant expression of mCherry in horizontal, amacrine, and ganglion cells (Figure 2C).

**Figure 2.**
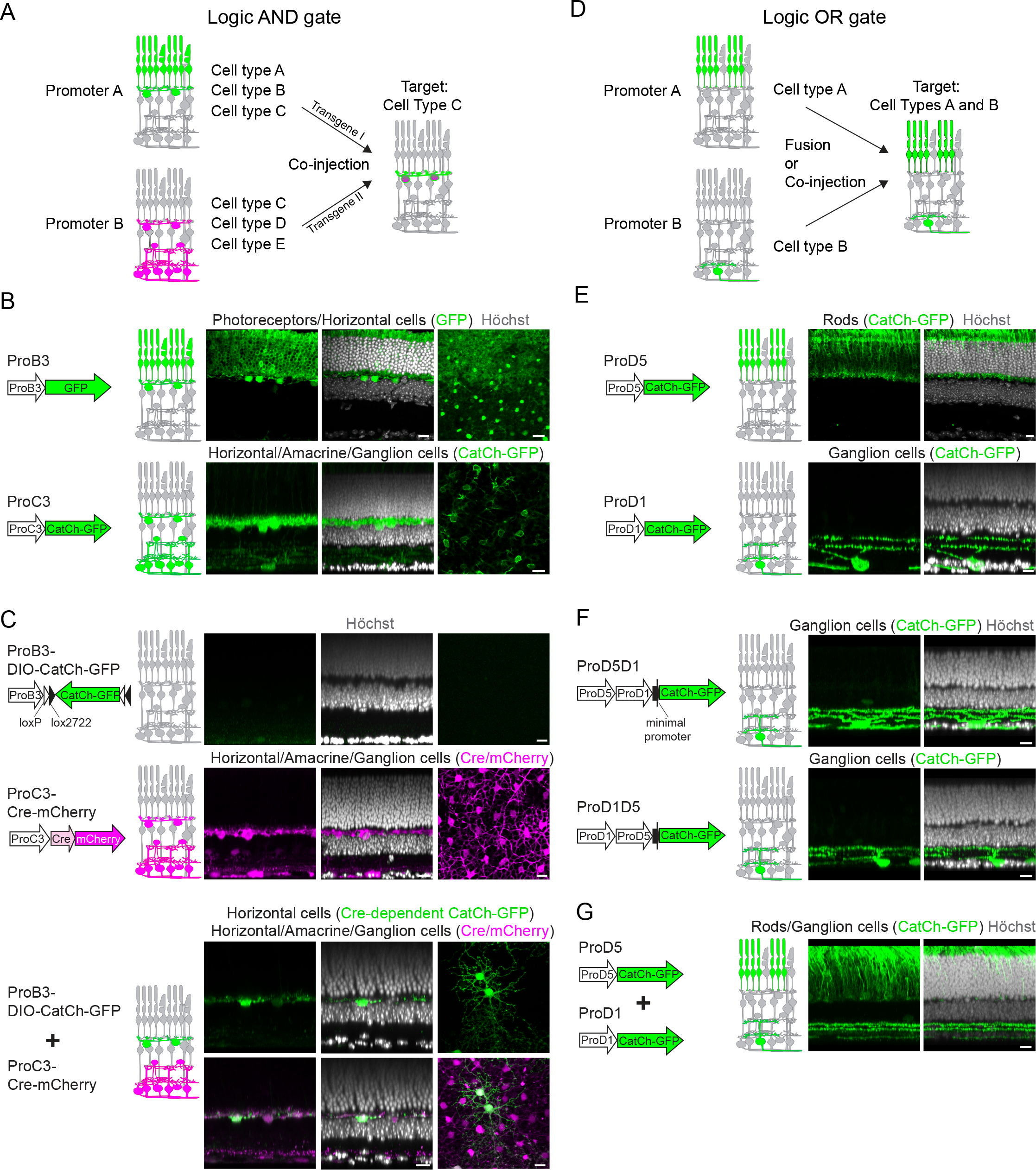
AND/OR logic for cell-type targeting by AAVs. (A) Logic AND gate strategy for cell-type targeting using two AAVs which themselves are not cell-type specific but which share one of their targets. (B-C) Confocal images of AAV-infected retinas using ProB3 or ProC3 synthetic promoters, which mutually target horizontal cells. Left and middle, retinal section; right, whole mount, confocal plane at the level of horizontal cells. Scale bars, 10 μm. (B) AAV-ProB3-GFP (top), AAV-ProC3-CatCh-GFP (bottom). (C) AAV-ProB3-DIO-CatCh-GFP (top), AAV-ProC3-Cre-mCherry (middle), coinjected Cre-dependent AAV-ProB3-DIO-CatCh-GFP and AAV-ProC3-Cre-mCherry (bottom). (D) Logic OR gate strategy for targeting selective combination of cell types using one or more AAVs. (E-G) Confocal images of AAV-infected retina sections using ProD5 and ProD1 synthetic promoters, which target rods and bistratified ganglion cells, respectively. Scale bars, 10 μm. (E) AAV-ProD5-CatCh-GFP (top), AAV-ProD1-CatCh-GFP (bottom). (F) AAV-ProD5-ProD1-CatCh-GFP (top), AAV-ProD1-ProD5-CatCh-GFP (bottom). (G) AAV-ProD5-CatCh-GFP and AAV-ProD1-CatCh-GFP co-injected.

Some applications requiring targeting of a particular combination of cell types may be carried out by using more than one synthetic promoter. We tested two strategies to combine synthetic promoters in order to express genes in a combination of cell types targeted by individual promoters (logic OR gate) (Figure 2D). First, we fused two synthetic promoters with differing cell-type specificities (Figure 2E) within a single AAV. Eyes were injected with an AAV carrying rod-targeting ProD5 fused to bistratified ganglion cell-targeting ProD1 and driving CatCh-GFP. In all AAV-infected retinas, CatCh-GFP was expressed exclusively in bistratified ganglion cells but not in rods, independent of the promoter structure, i.e., ProD5 first or ProD1 first (Figure 2F). The results suggested that fused sequences compromised ProD5 specificity and/or efficiency. Next, we co-injected eyes with a mixture of two different AAVs driving CatCh-GFP under ProD5 or ProD1. Dual AAV delivery efficiently targeted both rods and bistratified ganglion cells according to promoter specificity (Figure 2G). Thus, a combination of AAVs via AND/OR logic extends the repertoire of cell types that can be specifically targeted.

### Recording and modulating activity of AAV-targeted cells

AAV-mediated cell-type targeting creates an opportunity to record or modulate the functions of specific cell types. To test whether the extent of AAV-driven transgene expression is sufficient to monitor the activity of targeted cells, we performed *ex vivo* and *in vivo* Ca^2+^ imaging during visual stimulation. In retinas infected with AAV-ProA1 expressing a fluorescent Ca^2+^ indicator (GCaMP6s) in cones (Figure S2), two-photon imaging revealed a strong (>50% Δ*F/F*) light-evoked decrease in fluorescent traces in 93.8% of GCaMP6s-labeled cone terminals (Figure 3A), with a polarity typical of cone physiological light responses (Yau and Hardie, 2009). Next, we tested light-evoked Ca^2+^ transients in Müller glia cells. Glia cells sense and respond to neuronal activity through neuron-to-glia or glia-to-neuron signaling (Metea and Newman, 2006). In retinas infected with AAV-ProA18, which expresses GCaMP6s in Müller glia cells (Figure S2), two-photon imaging of cell terminals indicated a sustained increase in fluorescence in response to light (Figure 3B) that corresponded to a light-evoked increase in Ca^2+^ (Newman, 2005). The bistratified ganglion cells targeted by AAV-ProD1 (Figures 1C and S2) have the typical morphology of ON-OFF direction-selective (DS) ganglion cells. To test whether and in which direction ganglion cells highlighted by AAV-ProD1 are tuned, we infected retinas with AAV-ProD1-GCaMP6s, and analyzed cell responses following visual motion stimulation in eight different directions. Remarkably, all targeted ganglion cells showed vertical motion selectivity, with a dominating dorsally-tuned subtype (Figure 3C). We also analyzed the visual responses of AAV-targeted cells *in vivo* by injecting retinas with AAV-ProA5 to introduce GCaMP6 into ganglion cells (Figure S2). Neuronal activity of GCaMP6s-expressing ganglion cell axons was detected via two-photon imaging in the lateral geniculate nucleus (LGN). Light stimulation induced a significant fluorescence increase in a subset of axonal segments (Figure 3D). Altogether, these data demonstrate that AAV-mediated targeting allows monitoring of the activity of cells via GCaMP6s expression both *ex vivo* and *in vivo*.

**Figure 3.**
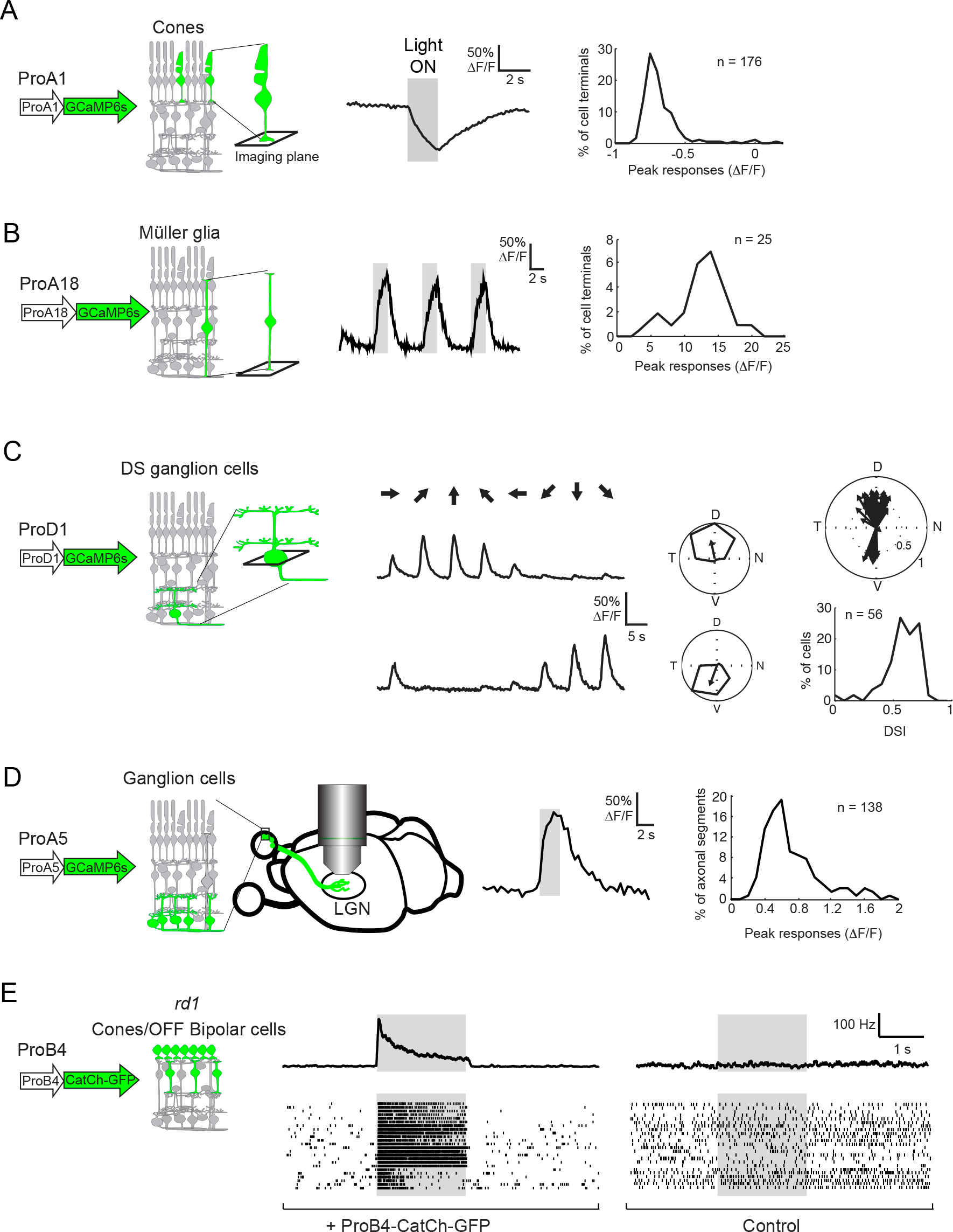
Recording and modulating activity of AAV-targeted cells. (A) Left, example of light-evoked decrease in fluorescence in ProA1-targeted cone terminals expressing GCaMP6s. Right, distribution of peak responses. (B) Left, example of light-evoked increase in fluorescence in ProA18-targeted Müller glia cell terminals expressing GCaMP6s. Right, distribution of peak responses. (C) Left, two examples (top and down) of visual-motion-induced responses in ProD1-targeted ganglion cells expressing GCaMP6s. Arrows indicate directions of visual motion. Middle, polar plots indicate response magnitudes of the recorded two cells normalized to the maximum response. Arrows indicate the vector sum of the responses; their direction is the preferred direction. Right, the preferred direction (arrow direction) and direction selectivity index (DSI; arrow length) of a set of ganglion cells targeted by AAV-ProD1 (top) and the distribution of their direction selectivity index (DSI, bottom). (D) Left, example of light-evoked increase in fluorescence in AAV-ProA5-targeted ganglion cell axon terminals in the lateral geniculate nucleus (LGN). Right, the distribution of peak responses. (E) Examples of light-evoked spike trains recorded in four ganglion cells in *rd1* retina infected with AAV-ProB4 targeting CatCh-GFP expression to OFF bipolar cells and cones (left) and in four ganglion cells in uninfected *rd1* retina (right). See also Figure S2.

We tested whether AAV-mediated cell-type targeting can modulate the activity of specific cell types. AAV-ProB4 was used to selectively target a type of retinal OFF bipolar cell, together with residual cones in *rd1* mice, a model of retinal degeneration lacking rod and cone responses by postnatal day 30 (P30) (Farber et al., 1994) (Figure S2 and Table S1). We examined whether optical stimulation of OFF bipolar cells/cones targeted with AAV-ProB4-CatCh-GFP evokes light responses in ganglion cells, measured as spike activity using a high-density multi-electrode array. CatCh-GFP activation by light stimulus led to both transient and sustained spike activity in cells in infected *rd1* retinas, but not in controls of the same age (>P30, Figure 3E). Thus, AAV-ProB4-induced expression of CatCh-GFP was sufficient to allow optogenetic stimulation.

### Cell-type targeting in mouse brain *in vivo*

We examined the use of the AAV library to target cell types in the mouse brain, choosing the visual cortex (V1) as a primary target. Four weeks after injection of 48 selected AAVs into V1 of adult mice, immunofluorescence analyses revealed induced transgene expression by 40% of the AAVs tested, with AAV-ProC17 and AAV-ProB12 highlighting the most restricted cell populations. To confirm and better characterize the targeted cells, we coated AAV-ProC17-CatCh-GFP and AAV-ProB12-CatCh-GFP with PHP.B capsid, which allows efficient transduction of mouse central nervous system cells after intravenous injection (Deverman et al., 2016).

Intravenous delivery of AAV-ProC17 induced CatCh-GFP expression in parvalbumin-positive (PV^+^) neurons in V1 (Figure 4A,B) and in other brain regions such as the barrel cortex (S1BF) and amygdala (CeA) (Figure 4C). Intravenous administration of AAV-ProB12 led to CatCh-GFP expression in glia-like cells in the central layers of V1 (Figure 4D). The morphology of targeted cells resembled protoplasmic astrocytes (Bushong et al., 2002; Kim et al., 2016) (Figure S3). AAV-ProB12 also labeled cells in the retrosplenial cortex (RS), S1BF and in dorsal thalamus (DT) with a morphology similar to those found in V1 (Figure 4D). To define the molecular identity of the targeted cells we made an AAV expressing GFP-tagged ribosomal protein L10a driven by the ProB12 synthetic promoter. The expression of GFPL10a enables translating RNAs from targeted cells to be isolated and analyzed (Nectow et al., 2017). We injected AAV-ProB12-GFPL10a intravenously, and isolated translating mRNAs from different brain regions. In the ProB12-GFPL10a^+^ cells, qRT-PCR of translating mRNAs revealed enrichment of astrocyte-specific transcripts (such as Gja1, Gjb6, Slc1a2, Slc1a3, Aqp4, Grm3, Aldoc, AI464131) (Lovatt et al., 2007; Cahoy et al., 2008; Zhang et al., 2014), astrocyte-enriched transcripts regulating cellular development (Sox9, Fgfr3, Tagln3, Mlc1) or metabolism (Cbs, Plcd4, Ppp1r3c), transcripts encoding astrocyte-enriched transmembrane receptors (Ntsr2, Gpr37l1, Ptprz1, Ednrb, S1pr1, Vcam1, Gria2, Dag1), ligands (Cpe, Mfge8, Htra1, Scg3, Pla2g7, Timp3, Btbd17, Itih3, Lcat), and secretion proteins (Clu, Cst3) (Lovatt et al., 2007; Doyle et al., 2008; Cahoy et al., 2008; Cordero-Llana et al., 2011) (Figure 4E). The ProB12-targeted cells did not express oligodenderocyte-(Plp1, Ndrg1, Mal, Omg, Mobp, Mbp, CNPase) or microglial-cell-specific markers (Tnf, C1qa, Ccl3, Iba1) (Sofroniew and Vinters, 2010; Watanabe et al., 2006; Barbarese et al., 1988) (Figures 4E and S3). Taken together, our data indicate that AAV-ProB12 targets a subpopulation of protoplasmic astrocytes. Within the cortex, this subpopulation is positioned in a central band.

**Figure 4.**
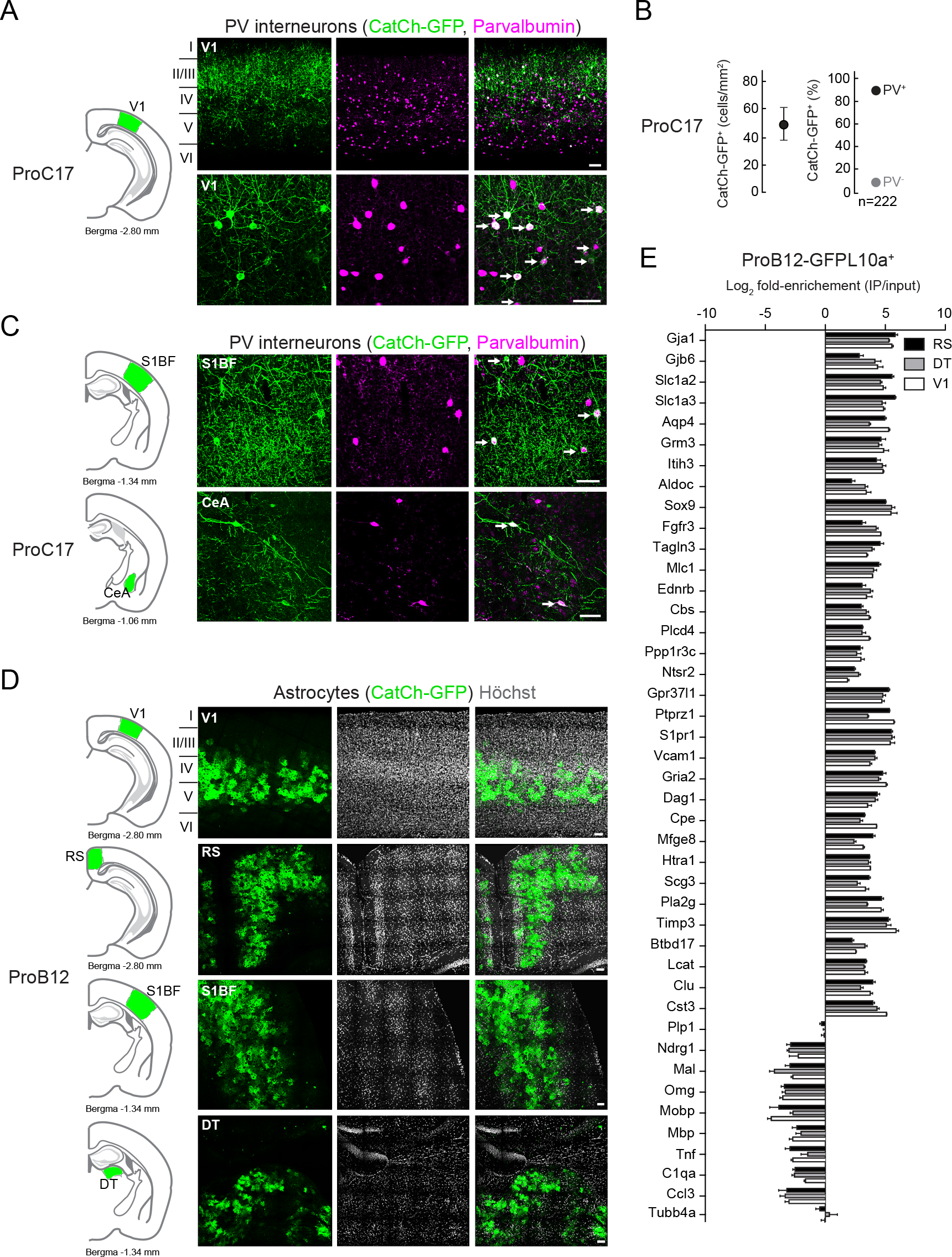
Cell-type targeting in mouse brain *in vivo*. (A) Confocal images of AAV-ProC17-infected sections of V1. Top, low magnification; bottom, high magnification. Left, CatCh-GFP (green); middle, immunostaining for PV (magenta); right, CatCh-GFP and PV. Arrows, co-labeled cells. Scale bars, 50 μm. (B) Left, quantification of CatCh-GFP^+^ cell density, mean ± SEM from three animals. Right, quantification of AAV-targeting specificity shown as a percentage of the PV (black) and non-PV (grey) cell types among cells expressing CatCh-GFP^+^. (C) Confocal images of AAV-ProC17-infected sections of two different brain regions, S1BF (top) and CeA (bottom). Left, CatCh-GFP (green); middle, immunostaining for PV (magenta); right, CatCh-GFP and PV. Arrows, co-labeled cells. Scale bars, 50 μm. (D) Confocal images of AAV-ProB12-infected sections of different brain regions. Top to bottom, V1, RS, S1BF, DT. Left, CatCh-GFP (green); middle, Höchst (white); right, CatCh-GFP and Höchst. Scale bars, 50 μm. (F) qRT-PCR quantification (mean ± SEM) of the set of translated mRNA expression in AAV-ProB12-GFPL10a-targeted cells isolated from the RS, DT, and V1 brain regions. Also see Figure S3.

### Cell-type targeting in non-human primate retina *in vivo*

To determine whether the AAVs generated target cell types in species other than mice, we tested the specificity of the AAVs in the retina of *Macaca fascicularis in vivo*. We injected eyes subretinally with 94 different AAVs, administrating four viruses with different synthetic promoters into four distinct quandrants of each eye. Analysis of transgene-expressing cells three months post-injection identified 62 AAVs that induced transgene expression. Photoreceptors were targeted individually or together with other cell types by 22 AAVs (Table S2). Five synthetic promoters, ProA1, ProA4, ProA7, ProB8, and ProD6, drove transgene expression specifically in cones, with ProA7 showing the highest targeting efficiency of 8,758±1,099 GFP^+^ cells per mm^2^. ProC1, ProC11, and ProD5 were rod specific, whereas eight others, including ProA6, targeted transgene expression to both photoreceptor types (Figures 5A,B and S4A and Table S2).

**Figure 5.**
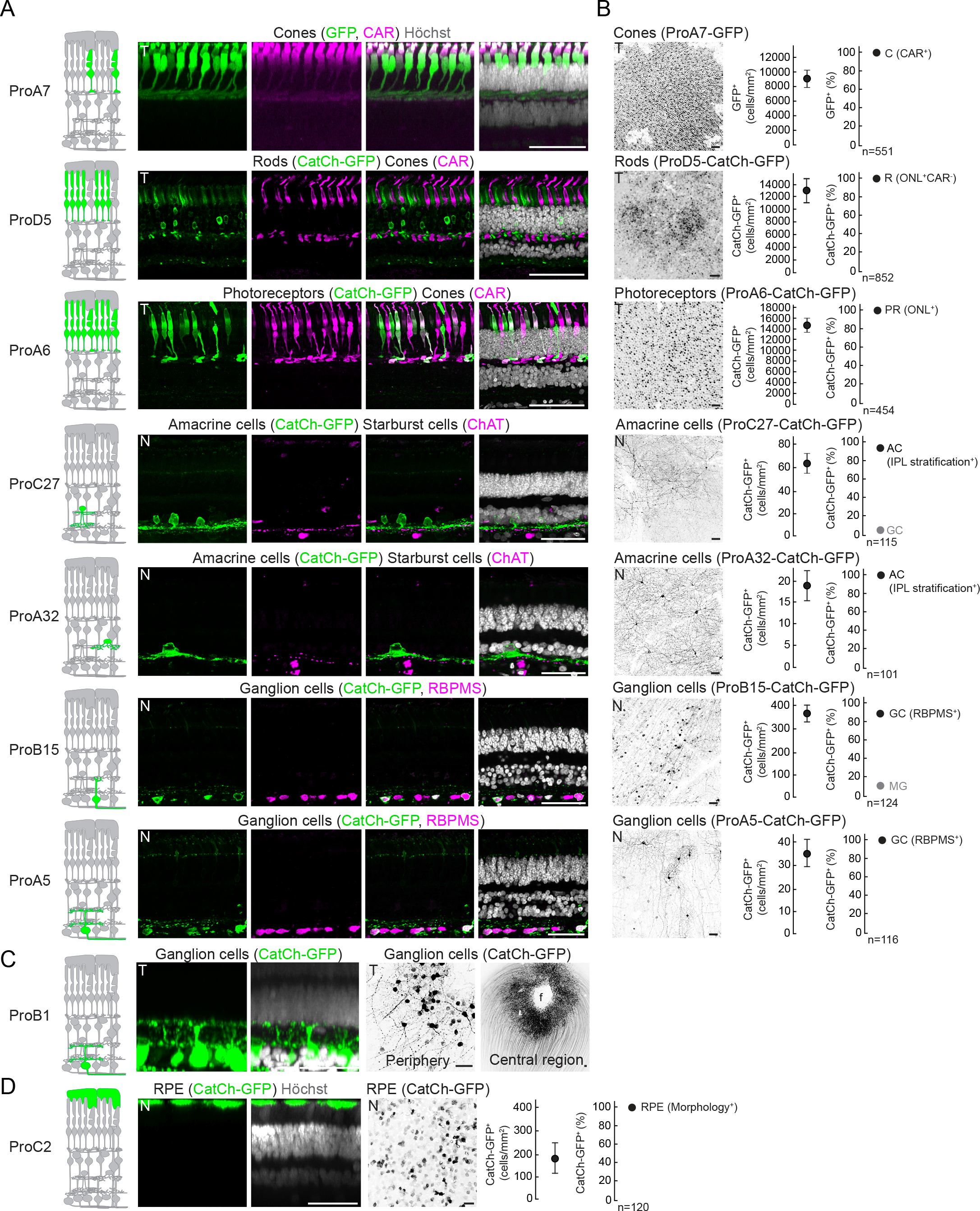
Cell-type targeting in non-human primate retina *in vivo.* (A) Confocal images of AAV-infected retinas. Left, GFP or CatCh-GFP (green); middle-left, immunostaining with marker (magenta) indicated above; middle-right, GFP or CatCh-GFP and marker; right, GFP or CatCh-GFP and marker and nuclear stain (Höchst, white). (B) Left, confocal images of AAV-infected retinas (top view), GFP or CatCh-GFP (black). Middle, quantification of GFP^+^ or CatCh-GFP^+^ cell density, values are means ± SEM from four retinas. Right, quantification of AAV-targeting specificity shown as a percentage of the major (black) and minor (grey) cell types among cells expressing the transgene. (C) Confocal images of AAV-ProB1-injected retina sections (left) and top views (right). Left, CatCh-GFP (green) and nuclear stain (Höchst, white). Right, CatCh-GFP (black) in peripheral retina and around the fovea (f). (D) Confocal images of a retina infected with AAV-ProC2. Left, retina sections showing CatCh-GFP (green) and nuclear stain (Höchst, white). Middle, confocal images of AAV-infected retinas (top view), CatCh-GFP (black). Right, quantification of CatCh-GFP^+^ cell density, values are means ± SEM from four retinas. Quantification of AAV-targeting specificity shown as a percentage of the major (black) cell types among cells expressing the transgene. C, cones; R, rods; PR, photoreceptors; MG, Müller glia; AC, amacrine cells; GC, ganglion cells; T, temporal retina quarter; N, nasal retina quarter. Scale bars, 50 μm. See also Figure S4 and Table S2.

Several synthetic promoters preferentially labeled different types of amacrine and ganglion cells (Figures 5 and S4 and Table S2), and promoters such as ProB15 and ProA5 targeted morphologically distinct ganglion cell types. Quantification of the dendritic-field diameter of AAV-targeted cells showed that AAV-ProB15 targeted cells with small and compact dendritic arbors (<100 μm), whereas ProA5 highlighted ganglion cells with larger cell bodies and dendritic fields (>100 μm) (Figures 5A,B and S4B). AAV-ProB1 highlighted a set of ganglion cells with restricted stratification in two IPL strata in the peripheral retina, as well as ganglion cells forming a circular rim around the fovea (Figure 5C). Three synthetic promoters (ProA18, ProB4, and ProC2) drove CatCh-GFP expression exclusively in the retinal pigmented epithelium (RPE), identified by the characteristic morphology and position of CatCh-GFP^+^ cells (Figures 5D and S4 and Table S2).

Taken together, our screen identified AAVs bearing a variety of synthetic promoters that preferentially targeted transgene expression to different cell types in the non-human primate retina.

### Cell-type targeting in the human retina *in vitro*

Using a culture protocol that we developed to keep *post mortem* human retinas alive for up to 14 weeks *in vitro*, we tested 84 AAVs, administering them on the peripheral retina either from the photoreceptor or the ganglion cell side. Immunofluorescence analyses after seven weeks revealed transduced individual or multiple cell types by 52 AAVs (Table S3). We identified two promoters (ProA7 and ProB8) that preferentially targeted cones, and six promoters, including ProA14, which co-targeted cones and rods (Figures 6 and S5 and Table S3).

**Figure 6.**
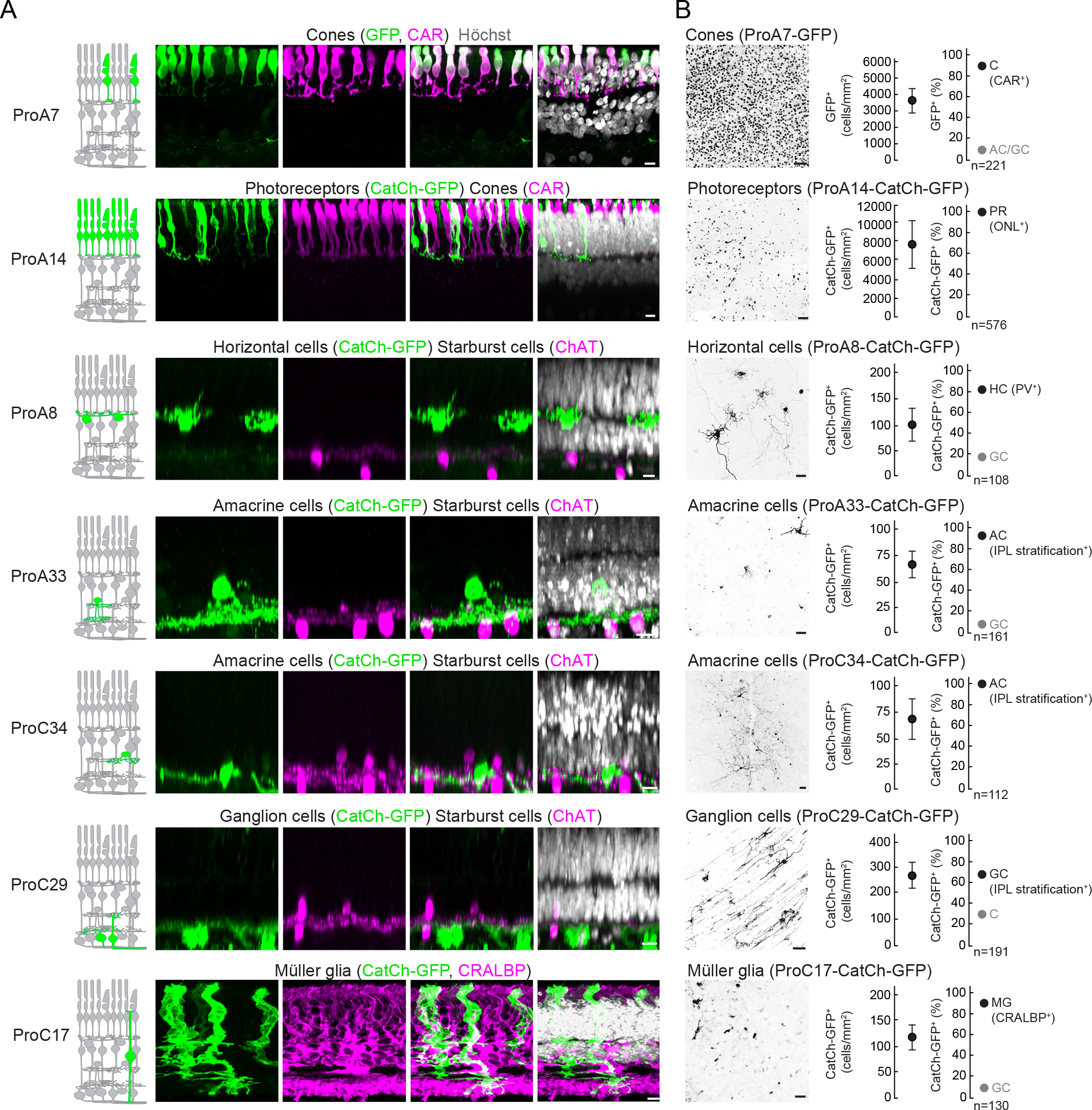
Cell-type targeting in human retina *in vitro.* (A) Confocal images of AAV-infected retinas. Left, GFP or CatCh-GFP (green); middle-left, immunostaining with marker (magenta) indicated above; middle-right, GFP or CatCh-GFP and marker; right, GFP or CatCh-GFP and marker and nuclear stain (Höchst, white). Scale bars, 10 μm. (B) Left, confocal images of AAV-infected retinas (top view), GFP or CatCh-GFP (black). Middle, quantification of GFP^+^ or CatCh-GFP^+^ cell density, values are means ± SEM from four retinas. Right, quantification of AAV-targeting specificity shown as a percentage of the major (black) and minor (grey) cell types among cells expressing the transgene. C, cones; R, rods; PR, photoreceptors; HC, horizontal cells; MG, Müller glia; AC, amacrine cells; GC, ganglion cells. Scale bars, 50 μm. See also Figure S5 and Table S3.

Human retinas contain three types of horizontal cells (Kolb et al., 1994). Horizontal cells are distinguished by the position of their cell body within the retina, their dendritic morphology, and co-labeling with a cell-type marker PV (Endo et al., 1986). Horizontal cell types differ in dendritic arbor size and branching pattern, connectivity to cone spectral subtypes, and in the lengths and terminal arborization of axons (Kolb et al., 1994). We found that AAVs such as ProA8, ProA29, and ProA37 sparsely targeted all three types of horizontal cells in the adult human retina (Figure 6 and Table S3). Considering inner retinal neurons, we found several synthetic promoters that highlighted different types of amacrine and ganglion cells (Figure 6). Few promoters induced CatCh-GFP expression preferentially in individual amacrine or ganglion cell types, but as many as 16 co-targeted mixed cell types from both classes (Table S3).

Finally, administration of AAV-ProC17 induced CatCh-GFP expression in Müller glia cells co-labeled with CRALBP (Figure 6).

### Correlation between targeting in mouse, non-human primate, and human retinas

For translational applications, it is useful to know whether a particular AAV targets the same cell types in humans as it does in mice or non-human primates. To reveal the relationship, we partitioned retinal cell types into eight cell groups (rods, cones, horizontal cells, bipolar cells, amacrine cells, ganglion cells, Müller glia, and retinal pigmented epithelium cells). For each AAV we created a vector with eight binary values: the value was 1 if expression was observed in a given cell group, and 0 otherwise. We then assessed the similarity of targeting across two species in two different ways: first, by computing the Pearson correlation (R) of the vectors between the two species (Figure 7A) and, second, by computing the conditional probability (CP) of expression in a given group in one species, given expression in the same group in a second species (Figure 7B).

**Figure 7.**
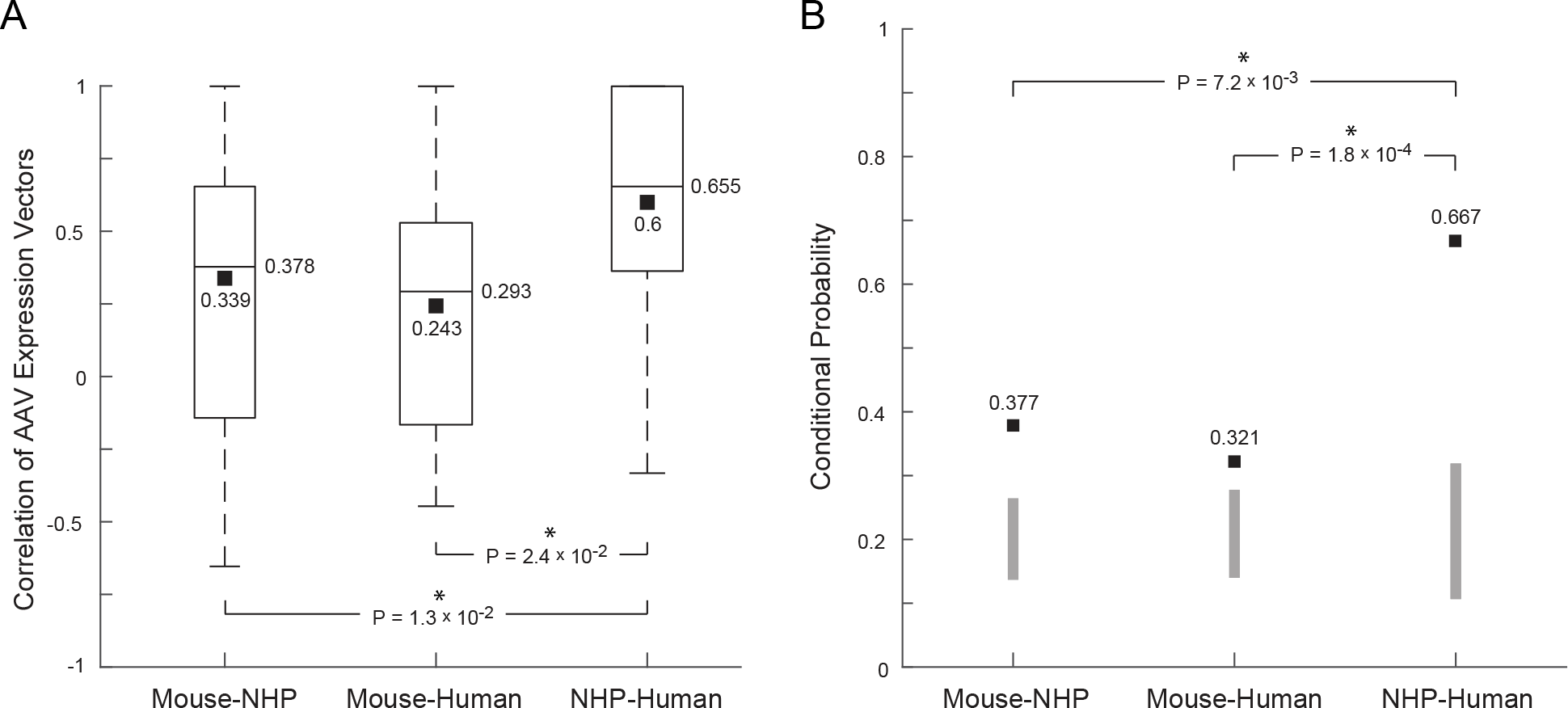
Quantitative metrics of the similarity of AAV expression in retinal cell groups in mice, non-human primates, and humans. (A) Box-and-whisker plots showing the distribution of the correlations between the AAV expression pattern vectors in two species: mouse and non-human primates (left), mouse and humans (middle), and non-human primates and humans (right). Boxes mark the 25^th^ and 75^th^ percentiles of the correlations together with the mean (black squares) and median (black lines). Statistical differences between correlation of AAV expression across pairs of species are shown numerically (stars indicate significance). (B) Mean conditional probability of AAV expression over all cell groups. Probability of expression in a cell group in one species, given expression in the same cell group in another (black squares) (solid, P < 0.05). The range of bars shows the distribution of conditional probabilities from a generative model of AAV expression assuming random expression in each species (mean ± 2 × standard deviation; generated by randomizing the AAV expression patterns 50,000 times). Statistical differences between different conditional probabilities are shown (stars indicate significance).

The mean correlation between mice and non-human primates, as well as between mice and humans, was significantly lower than the correlation between non-human primates and humans (mice/non-human primates, R = 0.339 ± 0.48; mice/humans, R = 0.243 ± 0.44; non-human primates/humans, R = 0.60 ± 0.50, mean ± standard deviation; P = 1.3 × 10^−2^, Monte Carlo sampling of difference distribution of correlations of mice/non-human primates and non-human primates/humans; P = 2.4 × 10^−2^, Monte Carlo sampling of difference distribution of correlations of mice/humans and non-human primates/humans; Figure 7A). Similarly, the conditional probability of observing expression in a cell group in non-human primates or humans, given that the same AAV expresses in that group in mice, independent of the specific cell group, was significantly lower than among non-human primates and humans (mice/non-human primates, CP = 0.377; mice/humans, CP = 0.321, non-human primates/humans, CP = 0.667; P = 7.2 × 10^−3^, Monte Carlo sampling of difference distribution of AAV expression in mice/non-human primates and non-human primates/humans; P = 1.8 × 10^−4^, Monte Carlo sampling of difference distribution of AAV expression in mice/humans and non-human primates/humans; Figure 7B). Nevertheless, all three conditional probabilities were significantly greater than predicted from randomizing the expression pattern of each AAV across the different cell groups (mice/non-human primates, P = 3.8 × 10^−8^; mice/humans, P = 1.3 × 10^−3^; non-human primates/humans, P = 9.5 × 10^−18^). Therefore, the ability of an AAV to target a cell group in mice is not a good predictor for targeting the same cell group in non-human primates and humans, although the association is not random. On the other hand, the ability of an AAV to target a cell group in non-human primates is a good predictor for targeting the same cell group in humans.

Since the expression pattern of AAV vectors in humans demonstrated a significantly greater conditional probability than expected from a randomly distributed expression pattern between mice and humans (Figure 7B), we asked whether experiments in mice could be helpful in restricting the number of AAVs to be further tested in non-human primates or humans. The conditional probabilities of an AAV expressing in a given cell group in non-human primates and humans, given that it does not express in the cell group in mice, are low (mice/non-human primates, CP = 0.12; mice/humans, CP = 0.14). Therefore it is reasonable to eliminate AAVs for testing in non-human primates and humans based on a lack of expression in mice.

## Discussion

Here we have described a collection of AAVs containing synthetic promoters that allow transgene expression in a broad range of retinal cell types in mice, non-human primates, and humans. Furthermore, using a subset of these AAVs, we present proof-of-concept for cell-type targeting with these synthetic promoters in other brain regions, such as the cortex. A few AAVs containing short regulatory sequences have been described previously that allowed neuronal and glial cell-type or broad cell-class targeting (Dimidschstein et al., 2016; Chaffiol et al., 2017; Lu et al., 2016; Cronin et al., 2014; Busskamp et al., 2010). However, there have been no reports to date of a broad spectrum of AAVs targeting neuronal or glial cell types, especially in non-human primates and humans. In combination with specific AAV serotypes and delivery routes, we have demonstrated the use of AAV-mediated cell-type targeting for morphological and molecular characterization as well as for monitoring and modulating the activity of specific cell types.

At the core of our AAV-based expression-targeting strategy is a regulatory DNA sequence, termed a synthetic promoter, embedded into the AAV vector genome 5’ from the gene to be expressed. Of the four different strategies for synthetic promoter design, none yielded sequences with a dominant cell-type-targeting specificity. The AAVs used here are episomal and do not integrate into the host cell genome (or only at a very low frequency) (Smith, 2008). It is likely that the molecular logic of the regulation of gene expression is different for episomal AAV DNA and host-cell DNA. The generation of effective synthetic AAV promoters likely depends on cell-or species-autonomous epigenetic and transcriptional regulatory mechanisms.

Importantly, the probability of an AAV targeting the same cell class in mice and non-human primates, or mice and humans, was as low as 0.38 and 0.32, respectively, compared with the probability for non-human primate and human retinal cell classes of 0.67. Therefore optimizing cell-type targeting in mice yields vectors that are unlikely to optimally target the same cell type in humans. However, our results do suggest that the absence of expression in a given cell type or cell class in mice is a useful proxy for the same in humans; therefore, studies in mice can be used to eliminate AAV vectors to be tested in humans.

The AAV resources described here have different potentials for basic and translational research in mice, non-human primates and humans.

In mice, AAVs make cell-type targeting fast and economical, and can thus benefit research. First, for cell-type targeting in mutant mice, since mating mice is costly and slow. Second, when the gene of interest changes often, such as optogenetic and activity sensor genes during improvement of these tools. Third, when analyzing connectivity across different neuronal cell types, for example by tagging one cell type with an optogenetic sensor and another with activity sensors; testing connectivity between the many possible cell-type pairs requires rapid and economic access to cell types. The ability to co-inject different AAVs to implement logic OR gates makes read-write connectivity mapping possible for cell-type pairs, and possibly also for n-tuples of cell types. The logic AND gates can help to target cell types for which no specific promoter is available or to induce sparse labeling. The route of administration of cell-type targeting AAVs determines whether a local subset of the cell type is labeled, or potentially the entire set. Local injection could result in the first scenario; systemic administration, using the PHP.B serotype for example, could result in the second scenario. An important consequence of screening cell-type-targeting AAVs is the discovery of new cell types. For example, in our screen in mice we discovered an unusual type of astrocyte. Within the cortex, this astrocyte type was restricted to the central layers.

In non-human primates, cell-type targeting by genome engineering is costly and time consuming and, thus, transgenic primates may be available for the study of only a few cell types. Therefore, for basic research in non-human primates, AAVs offer a simple, safe, and economic way to record or modulate the activity of cell types. Currently, only local labeling is available for these species. In translational research, AAVs can be used for *in vivo* proof-of-principle for gene therapy. Since non-human primate genetic disease models are increasingly available, it will become possible to demonstrate proof-of-concept for repair in the species closest to humans. This is important, since pre-clinical trials on mouse models often do not translate to humans, which is consistent with our finding that retinal cell-type targeting correlates well between non-human primates and humans, but much less well between mice and humans.

There is only limited knowledge about the functional roles of different cell types in the human brain. The culture of *post mortem* human brain parts, such as the retina or brain slices, in combination with cell-type targeting AAVs, could generate basic knowledge about, and understanding of, the organization and function of cell types in circuits in the human brain. Particularly notable for retinal research is the targeting of photoreceptors with optogenetic tools, and other cell types with genetically encoded activity sensors. Although the natural input from light is lost, restoring light sensitivity to photoreceptors may allow computations within the human retina to be studied for several months at the level of cell types and circuits. The human retina has a large surface area, 70 times larger than that of mice, and can be cut into many small retinas for independent studies. If cell-type targeting is achieved, all the tools used in mice will be available for human *post mortem* retinas, making it a simple and translationally relevant model system for research.

A prerequisite for human gene therapy is a vector system that allows efficient and long-lasting transgene expression in target cells. AAVs fulfill these criteria and are showing promise in clinical and pre-clinical gene therapy studies for inherited monogenic and complex eye diseases (Mendell et al., 2017; Russell et al., 2017). For example, AAV-based gene supplementation therapy has been successful in patients with Leber congenital amaurosis type 2, caused by a mutation in the RPE65 gene (Weleber et al., 2016; Russell et al., 2017). Gene supplementation cannot be used in the case of advanced retinal cell degeneration and, therefore, alternative strategies are needed to target and modulate the remaining retinal circuitry and to restore visual functions. AAVs in our collection that target human cones (e.g. ProA7) or ganglion cells (e.g. ProC29) could be used to express optogenetic tools that confer light-sensitivity on remnant cells in the diseased retina (Roska and Sahel, 2018). For both gene supplementation and optogenetic therapy, testing cell-type targeting in human retinas *in vitro* significantly increases the likelihood that the same vector will target the desired cell type in patients *in vivo*. This would provide a perspective for developing effective vectors for the treatment of blinding diseases such as Stargardt’s disease, age-related macular degeneration, Leber congenital amaurosis, retinitis pigmentosa, Leber hereditary optic neuropathy, dominant optic atrophy, and glaucoma. One drawback of testing AAVs in human retinas *in vitro* is that not all the factors are present, such as the immune system and the influence of surrounding tissues on AAV infection. Therefore, confirmation of expression from the same vectors in non-human primates is advisable before using them in clinical trials.

## Acknowledgments

We thank A.E. Kacso for multi-electrode array recording analyses, Z. Raics (SELS Software, Hungary) and D. Hillier for developing recording software, N. Ledergerber for assistance in mouse breeding and maintenance, A. Drinnenberg for providing AAV-ProA1-GCaMP6s confocal images, A. Police Reddy for assistance in cloning, X.W. Cheng for eye injections, L. Vandenberghe for advice on small-scale virus preparation, D. Gaidatzis for support in ProB synthetic promoters design, T. Siegmann for creating the AAV database, C. Cepko, V. Gradinaru, E. Bamberg, K. Deisseroth for providing plasmids and W. Baehr for providing the anti-CAR antibody. We thank P. King, S. Oakeley and E. Macé for commenting on the manuscript. We acknowledge the following grants: This work was supported by the Swiss National Science Foundation, the NCCR ‘Molecular Systems Engineering’, an ERC Advanced Grant, and a Gebert-Ruf grant to B.R., National Natural Science Foundation of China (81522014), National Key Research and Development Program of China (2017YFA0105300), and Zhejiang Provincial Natural Science Foundation of China (LQ17H120005) to Z.-B.J.

## METHODS

### CONTACT FOR REAGENT AND RESOURCE SHARING

Further information and requests for resources and reagents should be directed to and will be fulfilled by the lead contact, Botond Roska, upon signing of an MTA.

### EXPERIMENTAL MODEL AND SUBJECT DETAILS

#### Mice

Animals were used in accordance with standard ethical guidelines as stated in the European Communities Guidelines on the Care and Use of Laboratory Animals. C57BL/6J wild-type mice were obtained from The Charles River Laboratories. All mice were maintained in a pathogen-free environment with *ad libitum* access to food and drinking water. All animal experiments and procedures were approved by the Veterinary Department of the Canton of Basel-Stadt.

#### Non-human primates

Healthy cynomolgus monkeys (*Macaca fascicularis*, age 5-19 years, weight 5.6-10.8 kg) were housed at the Kunming Biomed International (KBI, Kunming, China) or Simian Laboratory Europe (Silabe, Strasbourg, France) facilities. The KBI is a facility accredited by the Association for Assessment and Accreditation of Laboratory Animal Care. Animals housed at the Silabe facility were maintained and monitored in accordance with the guidelines of the European Directive 2010/63, and handled in strict accordance with good animal practice as defined by the French National Charter on the Ethics of Animal Experimentation. All animal protocols were approved by the Institutional Animal Care and Use Committee of KBI or the French Ministry of Higher Education and Research (APAFIS#5716-2016061714424948v3).

#### Human retina tissue

Human retina tissue was collected from organ donors with no reported history of eye diseases at the Department of Ophthalmology, Semmelweis University (Budapest, Hungary). All tissue samples were obtained in accordance with the tenets of the Declaration of Helsinki. Personal identifiers were removed and samples were coded before processing. All experimental protocols were approved by the local ethics committees.

## METHOD DETAILS

### Synthetic promoter design

ProA synthetic promoters were based on mouse retinal cell type transcriptome profiling (Siegert et al., 2009; Hartl et al., 2017). The 2-kb nucleotide sequence upstream of the start codon of a highly cell-type-specific gene was selected as the synthetic promoter sequence. Most of the ProA synthetic promoters contain a short (<600 bp) region corresponding to the 5’ untranslated region of the dominant transcript isoform.

To generate the ProB synthetic promoters, highly conserved sequence elements were identified by whole vertebrate genome alignments using the UCSC genome browser. Elements that overlapped with repeat sequences were removed, and the conserved sequence elements were assigned to a gene by identifying the closest transcription start site. Conserved sequence elements far from a transcription start site were not considered or truncated in case of overlape with a transcript sequence. Conserved sequence elements were ordered according to the distance to the start codon of the closest cell-type-specific gene, or randomly assembled to generate the ProB synthetic promoter sequence.

ProC synthetic promoters were based on cell-type-specific transcription factor binding sites (Siegert et al., 2012) identified using the TRANSFAC and Jasper databases (Matys et al., 2003; Mathelier et al., 2016). Individual ProC synthetic promoters contained the selected transcription factor binding sites repeated 25 times interleaved by 15 bp random sequence spacers, followed by a TATA box containing an SV40 minimal promoter (Byrne et al., 1983).

ProD synthetic promoters are derived from *cis*-regulatory regions active in mouse retinal rods, cones, horizontal cells, and starburst amacrine cells that were identified using genome-wide DNA methylation maps (Hartl et al., 2017). The sequence of interest was PCR amplified from genomic mouse DNA (C57BL/6J) and supplemented with a synthetic minimal promoter sequence containing a TATA box (pGL.4.23-28, Promega), and universal primer binding and restriction sites (5’-atcctcacatggtcctgctggagttagtagagggtatataatggaagctcgacttccagctatcacatccactgtgttgttgtgaactggaatccactataggcca).

### AAV plasmid construction

Synthetic promoter sequences were chemically synthesized by Genewiz Inc. (South Plainfield, USA), with short flanks containing MluI/NheI/AscI and BamHI/EcoRI/BglII restriction sites. Synthetic promoter sequences were subcloned using an appropriate restriction site combination into pAAV-EF1a-CatCh-GFP or pAAV-hRO-GFP (Busskamp et al., 2010), replacing the EF1a or hRO promoters. The pAAV-EF1a-CatCh-GFP plasmid was constructed by adaptor PCR and the Clontech in-fusion kit using pcDNA3.1(-)-CatCh-GFP (a kind gift of E. Bamberg, MPI) and pAAV-EF1a-GFP (B. Roska lab plasmid collection). For Ca^2+^ imaging, ProA1, ProA5, ProA18, and ProD1 synthetic promoters were subcloned into pAAV-EF1a-GCaMP6s-WPRE-pGHpA via MluI/BamHI restriction sites replacing the EF1a promoter. For pAAV-ProB3-DIO-CatCh-GFP generation, the EF1a promoter was replaced with the ProB3 synthetic promoter using MluI/BamHI in the pAAV-EF1a-DIO-CatCh-GFP backbone (Roska lab plasmid collection). pAAV-ProC3-Cre/mCherry was generated using MluI/BamHI restriction sites replacing the EF1a promoter with ProC3 in the pAAV-EF1a-Cre/mCherry backbone (Roska lab plasmid collection).

### AAV production and titration

HEK293T cells were co-transfected with an AAV transgene plasmid, an AAV helper plasmid encoding the AAV Rep2 and Cap proteins for the selected capsid (8, 9, BP2, or PHP.B), and the pHGT1-Adeno1 helper plasmid harbouring adenoviral genes (kindly provided by C. Cepko, Harvard Medical School, Boston, USA) using branched polyethylenimine (PEI, Polysciences). For the “rapid AAV” protocol, one cell culture dish 15 cm in diameter was co-transfected with the plasmid mixture at 80% confluence of HEK293T cells. A cell transfection mixture containing 7 μg AAV transgene plasmid, 7 μg Rep2 and Cap-encoding plasmid, 20 μg AAV-helper plasmid, and 6.8 μM PEI in 5 ml of DMEM was incubated at room temperature for 15 min before being added to a cell culture dish containing 10 ml of DMEM. At 60 hrs post-transfection, cells were harvested and resuspended in buffer containing 150 mM NaCl and 20 mM Tris-HCl pH 8.0. Cells were lysed by repeated freeze-thaw cycles and MgCl_2_ was added to make a final concentration of 1mM. Plasmid and genomic DNA were removed by treatment with 250 U/ml of turbonuclease at 37ºC for 10 min. Cell debris was removed by centrifugation at 4,000 rpm for 30 min. AAV particles were purified and concentrated in Millipore Amicon 100K columns (Amicon, UFC910008, Millipore). Encapsidated viral DNA was quantified by TaqMan RT-PCR (forward primer: ggctgttgggcactgacaa; reverse primer: ccaaggaaaggacgatgatttc; probe: tccgtggtgttgtcg) following denaturation of the AAV particles using protease K, and titers were calculated as genome copies (GC) per ml. For the “conventional AAV” protocol, 10 cell culture dishes 15 cm in diameter were co-transfected with the mixture of AAV transgene, Rep2-Cap-encoding, and helper plasmids at 80% confluence of HEK293T cells. The AAVs were isolated using a discontinuous iodixanol gradient (OptiPrep, D1556, Sigma) and ultracentrifuged for 90 min at 242,000 g (Grieger et al., 2006b). AAV particles were purified and concentrated in Millipore Amicon 100K columns.

### AAV administration

Ocular injections were performed on mice anesthetized with 2.5% isoflurane. A small incision was made with a sharp 30-gauge needle in the sclera near the lens and 2 μl of AAV suspension was injected through this incision into the subretinal/intravitreal space using a blunt 5-μl Hamilton syringe held in a micromanipulator.

For intravenous administration, mice were anesthetized with 2.5% isoflurane and 30 μl of the AAV was injected into the retro-orbital vein using a 30-gauge micro-fine insulin syringe (Yardeni et al., 2011). A minimum of 3.3 × 10^11^ viral particles were injected per gram of mouse weight.

For non-human primate ocular injections, animals were anaesthetized with ketamine (10 mg/kg, Fujian Gutian Pharmaceutical Ltd, China) and phenobarbital sodium (5 mg/kg, Shanghai Xinya Pharmaceutical Ltd, China), and were positioned facing an operating microscope (66 Vision Tech Ltd, China). Pupils were dilated with 0.5% tropicamide / 0.5% phenylephrine hydrochloride (Santen, Finland). To visualize the fundus, a 30° circular prism (Suzhou MingRen, China) was placed on the cornea surface on top of medical sodium hyaluronate gel (Qisheng, China). Two 25-gauge pars plana sclerotomies were created and trocars were applied, enabling an illumination fiber (66 Vision Tech Ltd, China) and a 30-gauge needle mounted on a 50 μl Hamilton syringe (Hamilton, USA) to be inserted into the vitreous chamber. Different retinal quarters were subretinally injected with 50 μl of AAV (for titer specifications see Table S2).

### Human retinal culture

Human eyeballs were enucleated within 2 h of death under aseptic conditions, and rinsed in betadine (Egis Pharmaceuticals PLC, Hungary) for decontamination. The retina was dissected using fine scissors. For organotypic culture, 5 mm × 5 mm retinal pieces were isolated and placed ganglion cell or photoreceptor side up on polycarbonate membranes inserts (Corning). The culture were maintained at 37°C in 5% CO_2_ in DMEM/F12 medium (Thermo Fisher Scientific), supplemented with 0.1% BSA, 10 μM *O*-acetyl-L-carnitine hydrochloride, 1 mM fumaric acid, 0.5 mM galactose, 1 mM glucose, 0.5 mM glycine, 10 mM HEPES, 0.05 mM mannose, 13 mM sodium bicarbonate, 3 mM taurine, 0.1 mM putrescine dihydrochloride, 0.35 μM retinol, 0.3 μM retinyl acetate, 0.2 μM (+)-α-tocopherol, 0.5 mM ascorbic acid, 0.05 μM sodium selenite, 0.02 μM hydrocortisone, 0.02 μM progesterone, 1 μM insulin, 0.003 μM 3,3′,5-triiodo-L-thyronine, 2,000 U penicillin and 2 mg streptomycin (Sigma). For AAV infection, 20-40 μl of individual AAV (for titer specification see Tables S3) was applied per retina piece. AAV-induced transgene expression was examined 6-8 weeks after virus administration.

### Immunofluorescence and imaging

Retinas were fixed for 30 min in 4% (wt/vol) paraformaldehyde in PBS and washed with PBS for 24 h at 4°C. To improve antibody penetration, retinas were subjected to freeze/thaw cycles after cryoprotection with 30% (wt/vol) sucrose. After washing in PBS, retinal wholemounts or 3% agarose-embedded (SeaKem Le Agarose, Lonza) 150-μm-thick vibratome sections (Leica VT1000S vibratome) were incubated for 2 h in blocking buffer containing 10% (vol/vol) normal donkey serum (Chemicon), 1% (wt/vol) bovine serum albumin (BSA), 0.5% (vol/vol) TritonX-100, and 0.01% sodium azide (Sigma) in PBS. Primary antibody treatment was performed for 3-7 days at room temperature in buffer containing 3% (vol/vol) NDS, 1% (wt/vol) BSA, 0.01% (wt/vol) sodium azide, and 0.5% TritonX-100 in PBS. Primary antibodies used in this study: rabbit polyclonal anti-GFP (Thermo Fisher Scientific), rat monoclonal anti-GFP (Nacalai), chicken polyclonal anti-GFP (Thermo Fisher Scientific), rabbit polyclonal anti-mouse CAR (Millipore), mouse monoclonal anti-primate/human CAR 7G6 (Zhang et al., 2003), goat polyclonal anti-ChAT (Millipore), mouse monoclonal anti-CRALBP (Abcam), rabbit polyclonal anti-RFP (Rockland), guinea pig polyclonal anti-RBPMS (PhosphoSolutions), mouse monoclonal anti-mouse parvalbumin (Millipore), rabbit polyclonal anti-GFAP (Millipore), rabbit polyclonal anti-Iba1 (GeneTex), mouse monoclonal anti-CNPase (Millipore), rat monoclonal anti-MBP (Millipore), rat monoclonal anti-glycine (ImmunoSolutions), mouse monoclonal anti-tyrosine hydroxylase (TH) (Millipore). Secondary antibody incubation was performed for 2 h at room temperature in buffer supplemented with Höchst 33342 (10 μg/ml). Secondary antibodies used in this study: Alexa Fluor 488 donkey anti-rabbit IgG (H+L), Alexa Fluor 568 donkey anti-rabbit IgG (H+L), Alexa Fluor 647 donkey anti-rabbit IgG (H+L), Alexa Fluor 488 donkey anti-rat IgG (H+L), Alexa Fluor 488 donkey anti-mouse IgG (H+L), Alexa Fluor 555 donkey anti-mouse IgG (H+L), Alexa Fluor 647 donkey anti-mouse IgG (H+L), Alexa Fluor 488 donkey anti-goat IgG (H+L), Alexa Fluor 568 donkey anti-goat IgG (H+L), Alexa Fluor 633 donkey anti-goat IgG (H+L) (Thermo Fisher Scientific), Cy^™^3 AffiniPure donkey anti-rat IgG (H+L), Alexa488-conjugated donkey anti-chicken IgY, donkey anti-guinea pig Cy3 (Jackson Immuno Research). After washing in PBS, retinas were embedded in Prolong Gold antifade (Molecular Probes).

Brains were isolated and fixed for 2 h in 4% (wt/vol) paraformaldehyde in PBS and were then cut into 150-μm-thick slices using a vibratome. Slices were washed with PBS and incubated in blocking buffer followed by antibody staining as described above.

A Zeiss LSM 700 laser scanning confocal microscope was used to acquire images of antibody-stained retinas or brain slices with an EC Plan-Neofluar 40×/1.30 oil M27 and a Plan-Acro Achromat 10×/0.45 objective at up to four excitation laser lines according to secondary antibody specification. The morphologies of cell types were assessed from 512×512 pixel images in a z-stack with 0.85 μm z-steps. Images were processed using Imaris (Bitplane).

### Translated mRNA purification and qRT-PCR

Affinity purification of GFP-tagged polysomes from AAV-ProB12-GFPL10a-targeted cells was carried out 6 weeks after intravenous virus administration. Three biological replicates consisting of pooled brain regions isolated from two to three mice (mixed sex) were collected. Isolated brain regions were homogenized in buffer containing 10 mM HEPES-KOH (pH 7.4), 150 mM KCl, 5 mM MgCl_2_, 0.5 mM DTT, 100 μg/ml cycloheximide, RNasin and SUPERase-In RNase inhibitors, and Complete-EDTA-free protease inhibitors and then cleared by two-step centrifugation to isolate the polysome-containing cytoplasmic supernatant, as described previously (Heiman et al., 2014). Polysomes were immunoprecipitated using anti-GFP antibody bound to magnetic beads (GFP-Trap_MA, Chromotek), and bound RNA was purified using an Absolutely RNA Nanoprep kit (Agilent Technologies). RNA quantity and quality were measured on an Agilent 2100 Bioanalyzer. cDNA was prepared using a SensiFAST cDNA synthesis kit (BIOLINE, Switzerland). The SYBR Green method was used for qRT-PCR analysis on three biological replicates for each brain region, using a QuantStudio 3 real-time PCR system. Oligonucleotides used in this study (5՝ to 3՝ sequence): mmGja1F aacagtctgcctttcgctgt, mmGja1R atcttcaccttgccgtgttc, mmGjb6F caggtttgggtgttttgctt, mmGjb6R ctcatcaccccacacttcct, mmSlc1a2F agagggtgccaacaatatgc, mmSlc1a2R atgaccacatcagggtggat, mmSlc1a3F ccaaaagcaacggagaagag, mmSlc1a3R acctcccggtagctcatttt, mmAqp4F agcaattggattttccgttg, mmAqp4R tgagctccacatcaggacag, mmGrm3F ggaaacattggacccactca, mmGrm3R caggcgttggatacctctgt, mmItih3F caagacagccttcatcacca, mmItih3R gttgtttgggctggacctta, mmAldocF aggcatcaaggttgacaagg, mmAldocR ataggcacgatcccattctg, mmSox9F tgcagcacaagaaagaccac, mmSox9R ccctctcgcttcagatcaac, mmFgfr3F cgtgtaacaggggctcctta, mmFgfr3R gtgtgtatgtctgccggatg, mmTagln3F tgagcaaattggtgaacagc, mmTagln3R ttccatcgtttttggtcaca, mmMlc1F ctgactcaaagcccaaggac, mmMlc1R gtagtcacagcgaacgtgga, mmEdnrbF atgacgccacccactaagac, mmEdnrbR gatgatgcctagcacgaaca, mmCbsF tggaaacttgaagcctggag, mmCbsR gcggtactggtccagaatgt, mmPlcd4F caagggttcaccgttgtctt, mmPlcd4R tccacgttcatcagaagcag, mmPpp1r3cF gtgaccgggacagtgaaagt, mmPpp1r3cR cactccgtcaggtttccatt, mmNtsr2F cgcctgctgtcactagtctg, mmNtsr2R gagttgacttgggcagaagc, mmGpr37l1F gagagctcctacagcgccta, mmGpr37l1R aacatccccgagtagccttt, mmPtprz1F attggctggtcctacacagg, mmPtprz1F tgcttgccttgaagaccttt, mmS1pr1F ctctgctcctgctttccatc, mmS1pr1R gatgatggggttggtacctg, mmVcam1F cccgtcattgaggatattgg, mmVcam1R taaggtgagggtggcatttc, mmGria2F atttcgggtagggatggttc, mmGria2R gttgggaagcttggtgtgat, mmDag1F gtgagcattccaacggattt, mmDag1R tggctcattgtggtcttcag, mmCpeF cgccatcagcagaatctaca, mmCpeR ggttcaaggagggcatgata, mmMfge8F ttctgtgactccagcctgtg, mmMfge8R tggcagatgtattcggtgaa, mmHtra1F acgccaagacctacaccaac, mmHtra1R tcctccgatacgatgaatcc, mmScg3F gccaccaggatttatgagga, mmScg3R ttttcccatcctgattctcg, mmPla2g7F tcggttatgggaatgagagc, mmPla2g7R attagatgccaagccaatgc, mmBtbd17F cagcttctgggctattctgg, mmBtbd17R cgcagcataacctcactctg, mmLcatF cttcaccatctggctggatt, mmLcatR ccagattctgcaccagtgtg, mmCluF tgagctccaagaactgtcca, mmCluR tcatgcaggtatgcttcagg, mmCst3F tcgctgtgagcgagtacaac, mmCst3R tgcagctgaattttgtcagg, mmPlp1F ctggctgagggcttctacac, mmPlp1R gactgacaggtggtccaggt, mmTimp3F cacggaagcctctgaaagtc, mmTimp3R cccaaaattggagagcatgt, mmNdrg1F aaccgtcctgtcatcctcac, mmNdrg1R tgccaatgacactcttgagc, mmMalF tccctgacttgctcttcgtt, mmMalR ggtaaaatagggcagccaca, mmOmgF ttggacagttccaaccaaca, mmOmgR cctatcacatgggctttcgt, mmMobpF tcaaccccaaggaagaagtg, mmMobpR tgaaaccaaaagacccgttc, mmMogF aaatggcaaggaccaagatg, mmMogR gacctgcaggaggatcgtag, mmMbpF gcttctttagcggtgacagg, mmMbpR gaggtggtgttcgaggtgtc, mmTnfF ccgatgggttgtaccttgtc, mmTnfR cggactccgcaaagtctaag, mmC1qaF acaaggtcctcaccaaccag, mmC1qaR tccttttcgatccacacctc, mmCcl3F accatgacactctgcaacca, mmCcl3R cccaggtctctttggagtca, mmTubb4aF agttagtggatgccgtcctg, mmTubb4aR ccagctgatgcacagacagt, mmTuba4aF tcagtgcacaggacttcagg, mmTuba4aR cggcggcagatatcataaat. Data were normalized to *Tuba4a* using the comparative C_T_ (2^−ΔΔCT^) method.

### Two-photon calcium imaging

Retinas infected with AAV-ProA1-, AAV-ProA18-, and AAV-ProD1-GCaMP6s were isolated and the pigment epithelium removed in Ringer’s solution (110 mM NaCl, 2.5 mM KCl, 1 mM CaCl_2_, 1.6 mM MgCl_2_, 10 mM D-glucose, 22 mM NaHCO_3_, bubbled with 5% CO_2_/95% O_2_, pH 7.4) and mounted ganglion-cell-side up on a filter MF-membrane (Millipore) with a 2-mm rectangular aperture in the centre. The retinas were superfused in Ringer’s solution at 35-36°C in the microscope chamber for the duration of the experiment. The two-photon microscope system has been described previously (Yonehara et al., 2013). Briefly, the system was equipped with a Mai Tai HP two-photon laser tuned to 920 nm (Spectra Physics) and a 60× objective (Fluor, 1.0 NA, Nikon). Image data were acquired using custom software developed by Z. Raics (SELS Software, Hungary), taking images of 150 × 150 pixels (10 frames per second, for ProA1 and ProA18) or 200 × 200 pixels (5.7 frames per second, for ProD1). A TTL signal generated at the end of each line scan of the horizontal scanning mirror was used to trigger a UV LED projector (Acer) (Reiff et al., 2010). To prevent stimulation light bleeding through and masking light emission from the sample, stimuli were presented exclusively during the fly-back period of the horizontal scanning mirror. Visual stimulation was generated via custom-made software (Python, Labview, National Instruments). The light intensity of the visual stimulation was 7.2 × 10^4^ photoisomerizations per rod per second (R*/s) with a background intensity of 1.4 × 10^2^ R*/s. For AAV-ProA1 and ProA18, the stimulation was a flash of a circular light spot of 120 μm diameter presented for two seconds. For AAV-ProD1, the stimuli were circular light spots of 400 μm moving at a speed of 800 μm/s on the retina in eight different directions.

The fluorescence data were analyzed semi-online via custom-made software written in Python by Z. Raics (SELS Software, Hungary). For ProA1, the cone axon terminals were automatically segmented via an algorithm developed by D. Hillier (Hillier et al., 2017). For ProA18 and ProD1, the terminal areas and the cell bodies were segmented manually. Background fluorescence was calculated as the mean of the 10% dimmest pixels for each frame, and subtracted from the mean fluorescence of each segmented area. The resulting fluorescence values were then normalized as ΔF/F, where F represents baseline (mean fluorescence of a 1-2 s time window before the onset of the stimulus). In the case of repetitive stimulation, all responses to different trials were averaged before calculating the peak response. Peak responses (ProA1 and ProA18) and preferred direction and direction-selective index (DSI, ProD1) were analyzed offline using MATLAB (MathWorks). Peak responses in cone axon terminals (ProA1) were calculated as the means of ΔF/F values during the second half of the stimulation period. Peak responses in Müller glia cells (ProA18) were the point of maximum fluorescence in the stimulation period. Preferred direction and DSI (ProD1) were calculated as previously described (Wertz. et. al., 2015). Briefly, for the cell of interest, eight vectors were formed first, each associated with motion along a different direction. The angle of the vector was the angle corresponding to the motion direction (0°, 45°, 90°, 135°, 180°, 225°, 270°, 315°). The length of each vector was the response amplitude along the relevant direction, where the amplitude was defined as the maximum ΔF/F in the peak. The preferred direction and DSI of the cell of interest were calculated as the angle and length of the sum of the eight vectors divided by the sum of the lengths of the eight vectors, respectively. Cells that showed no significant responses (signal-to-noise-ratio < 50, where signal-to-noise-ratio is defined as the amplitude of the largest peak divided by the standard deviation of the baseline) were excluded from the analysis. Finally, response histograms were assembled and plotted (Figure 3A,B,D).

### Two-photon imaging of ganglion cell axons in the LGN

AAVBP2-ProA5-GCaMP6s was administered into the right mouse retina as described above. After 3-4 weeks, mice were anesthetized with fentanyl/medetomidine/midazolam (fentanyl 0.05 mg/kg, medetomidine 0.5 mg/kg, midazolam 5.0 mg/kg) and placed in a stereotaxic frame (Narishige, SR-5M). Coliquifilm (Allergan) was applied to the eyes to prevent dehydration during surgery. A metal bar for head fixation during imaging was glued to the skull (Holtmaat et al., 2009). A 3-mm diameter craniotomy was made above the LGN. The exposed cortex and the underlying hippocampus were aspirated, exposing the LGN. The tissue was kept moist with mouse Ringer’s solution (135 mM NaCl, 5.4 mM KCl, 5 mM HEPES, 1.8 mM CaCl_2_, pH 7.2) heated to 37 ºC. A 2-mm diameter glass coverslip was slightly pushed against the LGN, while the tissue between the edges of the coverslip and the skull were covered with superglue and allowed to solidify. After surgery, anesthetized mice were placed under a two-photon microscope. Retinal ganglion cell axons in the left LGN were imaged through a 40× objective (LUMPlanFl 40×/0.8NA water immersion, Olympus) between 20 and 50 μm from the surface of the LGN. The mouse was presented with six flashes of blue light, each lasting 5 s, with 10 s between each light flash. Blue light was produced by a mounted LED (Thorlabs, M405L3) focused with an achromatic doublet lens (Thorlabs, AC254-030-A-ML) onto a fiberoptic cable (Thorlabs, M58L005), which was placed in front of the mouse to project full-field light onto the right eye. GCaMP6s responses were collected at 3 Hz (National Instruments, Labview) and analyzed in Matlab (MathWorks). Each retinal ganglion cell axon segment was selected manually and ΔF/F was calculated by dividing the average pixel intensity within the region of interest (ROI) for each timepoint by the mean signal intensity for that ROI during the 5 s before visual stimulation. An average light response trace was obtained by averaging the responses over the six light flashes. Peak responses were calculated from the maximal ΔF/F of average light responses in each axon segment during the 5 s of light stimulation.

### Multi-electrode array recordings

To record the spike trains of retinal ganglion cells, the retina of a wild-type C57BL/6J mouse or a mutant rd1 mouse infected with AAV-ProB4-CatCh-GFP was isolated under dim red light in Ringer’s solution (110 mM NaCl, 2.5 mM KCl, 1 mM CaCl_2_, 1.6 mM MgCl_2_, 10 mM D-glucose, 22 mM NaHCO_3_) bubbled with 5% CO_2_/95% O_2_. The retina, ganglion-side down, was then immobilized on the multi-electrode array by gently pressing with a cell culture membrane (Transwell 3450-Clear) bearing hexagonally arranged holes of 200 μm diameter and a center-to-center distance of 400 μm. For the duration of the experiment, the retina was perfused with oxygenated Ringer’s solution (110 mM NaCl, 2.5 mM KCl, 1.0 mM CaCl_2_, 1.6 mM MgCl_2_, 22 mM NaHCO_3_, 10 mM D-glucose, pH 7.4, with 5% CO_2_/95% O_2_) at a flow rate of 1.5 ml/min at 35°C. Extracellular voltage was measured with a multi-electrode array (MEA1060 Up-BC amplifier, Multichannel Systems) at 20 kHz. The array was fixed onto a motorized table (Scientifica). CatCh was activated by a light stimulus (7 × 10^16^ photons/cm^2^/s) generated using a DLP projector (PLUS U137SF) and projected onto the retina by the condenser lens of an inverted microscope (Nikon TE300). The spectrum of the stimulus light was determined using a spectrophotometer (Ocean Optics USB2000) and the light intensity was measured using a power meter (Thorlabs S130VC). The stimulation intensity was calculated by integrating the product of the projector spectrum and the normalized absorption spectrum of CatCh-GFP. The recorded voltage was bandpass-filtered (400-4000 Hz) and spikes were sorted using the UltraMegaSort software (Kleinfeld Lab, University of California, San Diego). Spike frequency was calculated using 50-ms moving bins. Intrinsically photosensitive retinal ganglion cells were discriminated by their delayed spiking. For quantification, only spike frequency values in the first 200 ms after light start or end were used.

## QUANTIFICATION AND STATISTICAL ANALYSIS

### Correlation

The expression pattern of each AAV was divided into eight cell classes (rods, cones, horizontal cells, bipolar cells, amacrine cells, ganglion cells, Müller glia, retinal pigmented epithelium cells). Ignoring the relative penetrance of each AAV, the expression in each class was binarized. We computed the Pearson correlation of the expression vector in each pair of species for every AAV tested in both species. The reported results are the first-order statistics (mean, median; standard deviation, median) over the set of correlations for all AAVs in each pair of species. The correlation is a metric ranging in value from −1 to 1; if both expression pattern vectors are identical, the correlation is 1; if the expression in both species in every cell class is different, the correlation is −1.

### Monte Carlo sampling of difference distribution of correlations

We computed a statistical significance of the difference in the mean Pearson correlation between two different pairs of species, by using the two vectors of underlying correlations (each composed of the Pearson correlations of all the AAVs tested in each pair of species). We concatenated both vectors and resampled two novel vectors (of the same size as each original vector). We generated a novel random variable that was the difference between the mean of these two new vectors, sampling from a novel distribution which we term the *difference distribution*. Given H_0_, that the two distributions are statistically identical, this random variable will be normally distributed around 0. We repeatedly sampled from the difference distribution, computing all possible samples (up to a maximum of 50,000) and then computed the mean and standard deviation of the distribution. The true difference between the mean correlations between two pairs of species can be viewed as a potential sample from the difference distribution. Computing its *Z* score relative to the distribution then allowed us to compute the bounding probability that it is actually a sample from the difference distribution, which we have reported in the text.

### Conditional probability

As before, the expression pattern of each AAV was divided into eight retinal cell classes and appropriately binarized. For each pair of species, we counted the number of instances where an AAV was expressed in the same cell class in both species, independent of cell class. We then normalized this by the total number of times an AAV was expressed in each cell class in the first species (e.g. expression in mouse in the mouse/human pair), again independent of cell class. This gives an estimate of the probability that an AAV will be expressed in a cell class, given that it is expressed in the same cell class in one species, independent of the specific cell class. For the conditional probability given the lack of expression, we followed an analogous process.

### Monte Carlo sampling of difference distribution of AAV expression

We have reported two statistical tests of significance for the conditional probability. (1) The probability that the observed conditional probability of expression was actually drawn, in each species, independently from a random distribution. We generated this artificial distribution by randomly permuting the cell class in which each AAV was expressed, controlling for the number of cell classes in which it was expressed. We then generated 50,000 such randomized samples of AAV expression, in each of the two species. For each example, we computed the mean conditional probability between the two species, generating a sample from the distribution of conditional probability assuming that the expression of each AAV is independent between species. Since this distribution may not be normal, we computed the significance as introduced in the previous section, through the use of a difference distribution. In this case, the two vectors are of unequal size: the first contains the single observed value, and the second contains the 50,000 samples from the distribution derived from randomizing AAV expression. Computing the *Z* score of the difference between the observed value and the sampled distribution provided us with the probability that the observed conditional probability is actually sampled from a random distribution, which we have reported. (2) The probability that the observed conditional probability of expression in one pair of species is significantly different from the same in a second pair of species. Again, making no assumption about the underlying distribution, we utilized the same approach of generating a difference distribution. For each pair of species, we generated a vector of the conditional probability, by normalizing the number of cell classes where an AAV was expressed in the same cell class in both species by the total number of cell classes where the AAV was expressed in the first species. Then, using these two conditional probability vectors, we computed the difference distribution. As before, we sampled the difference distribution and use the *Z* score of the observed difference in conditional probability between the two pairs of species to compute the bounding probability that the two observed samples are actually from the same underlying distribution.

---

**Figure S1 related to Figure 1.**
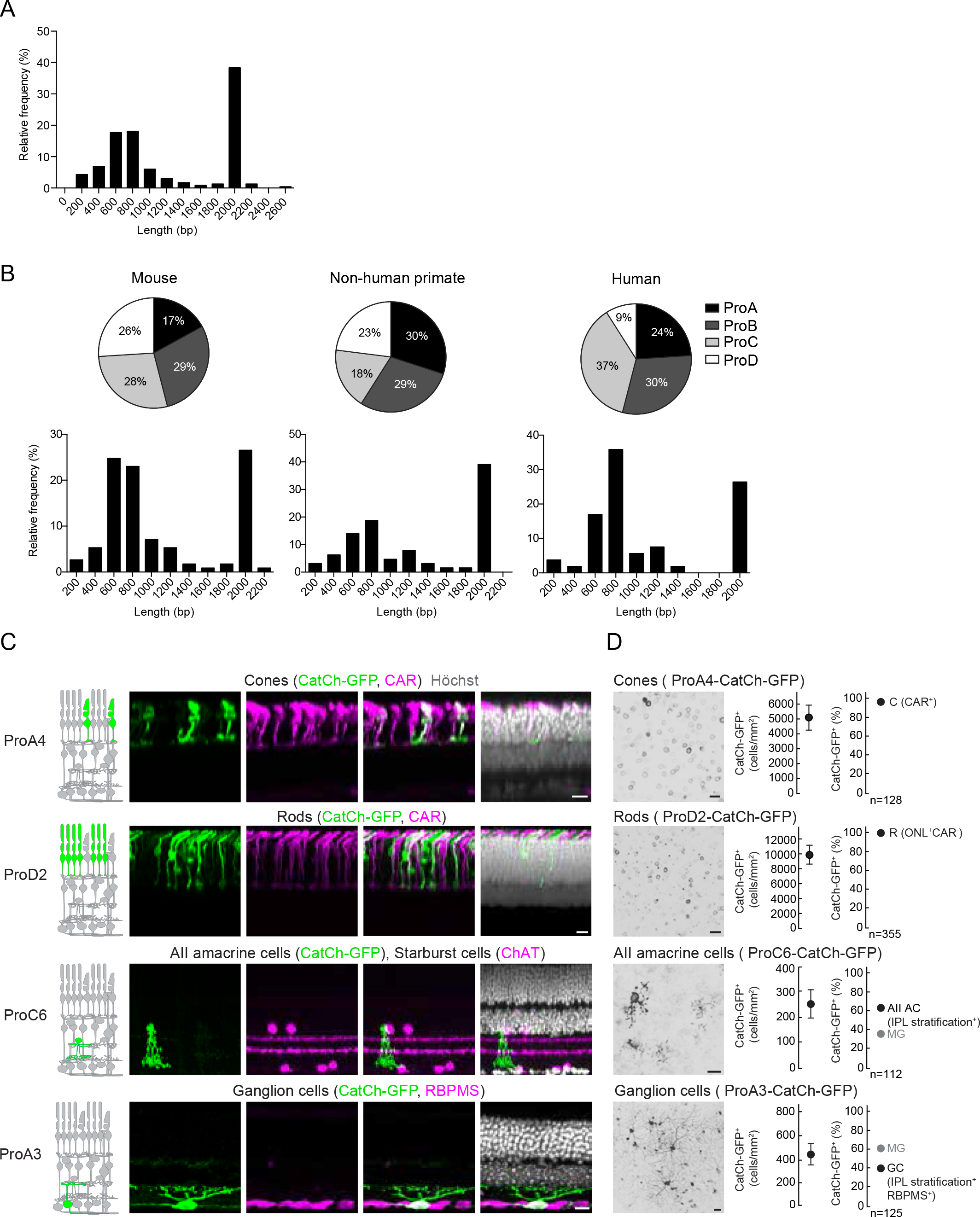
AAV-mediated cell-type targeting in mouse, non-human primate, and human retina. (A) Histogram of the relative frequencies of the lengths of synthetic promoters. (B) Pie charts (top) show percentages of each synthetic promoter group among synthetic promoters targeting transgene expression to at least one cell type in the mouse (left), non-human primates (middle), and humans (right) normalized to the number of promoters tested from each group. Histograms (bottom) show relative frequencies of active synthetic promoter lengths in different species. (C) Confocal images of AAV-infected retinas. Left, CatCh-GFP (green); middle-left, immunostaining with marker (magenta) indicated above; middle-right, CatCh-GFP and marker; right, CatCh-GFP and marker and nuclear stain (Höchst, white). Scale bars, 10μm. (D) Left, confocal images of AAV-infected retinas (top view), CatCh-GFP (black). Middle, quantification of CatCh-GFP^+^ cell density, values are means ± SEM from four retinas. Right, quantification of AAV-targeting specificity shown as a percentage of the major (black) and minor (grey) cell types among cells expressing the transgene. C, cones; R, rods; AII AC, AII amacrine cells; MG, Müller glia; GC, ganglion cells. Scale bars, 50 μm.

**Figure S2 related to Figure 3.**
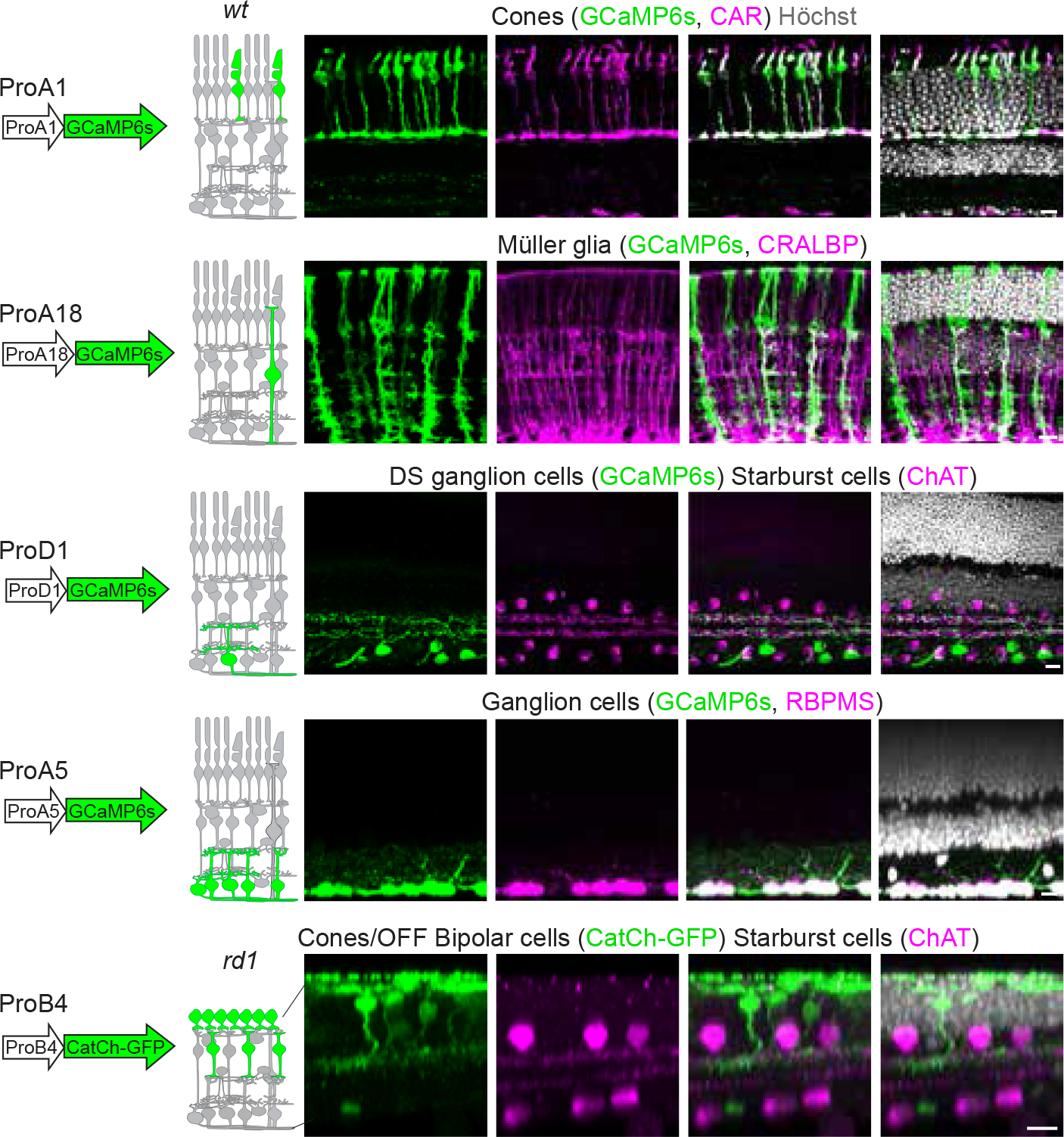
AAV-mediated GCaMP6s or CatCh-GFP expression in wild-type or *rd1* retinas. Confocal images of AAV-infected retinas. Left, GCaMP6s or CatCh-GFP (green); middle-left, immunostaining with marker (magenta) indicated above; middle-right, GCaMP6s or CatCh-GFP and marker; right, GCaMP6s or CatCh-GFP and marker and nuclear stain (Höchst, white). Scale bars, 10 μm.

**Figure S3 related to Figure 4.**
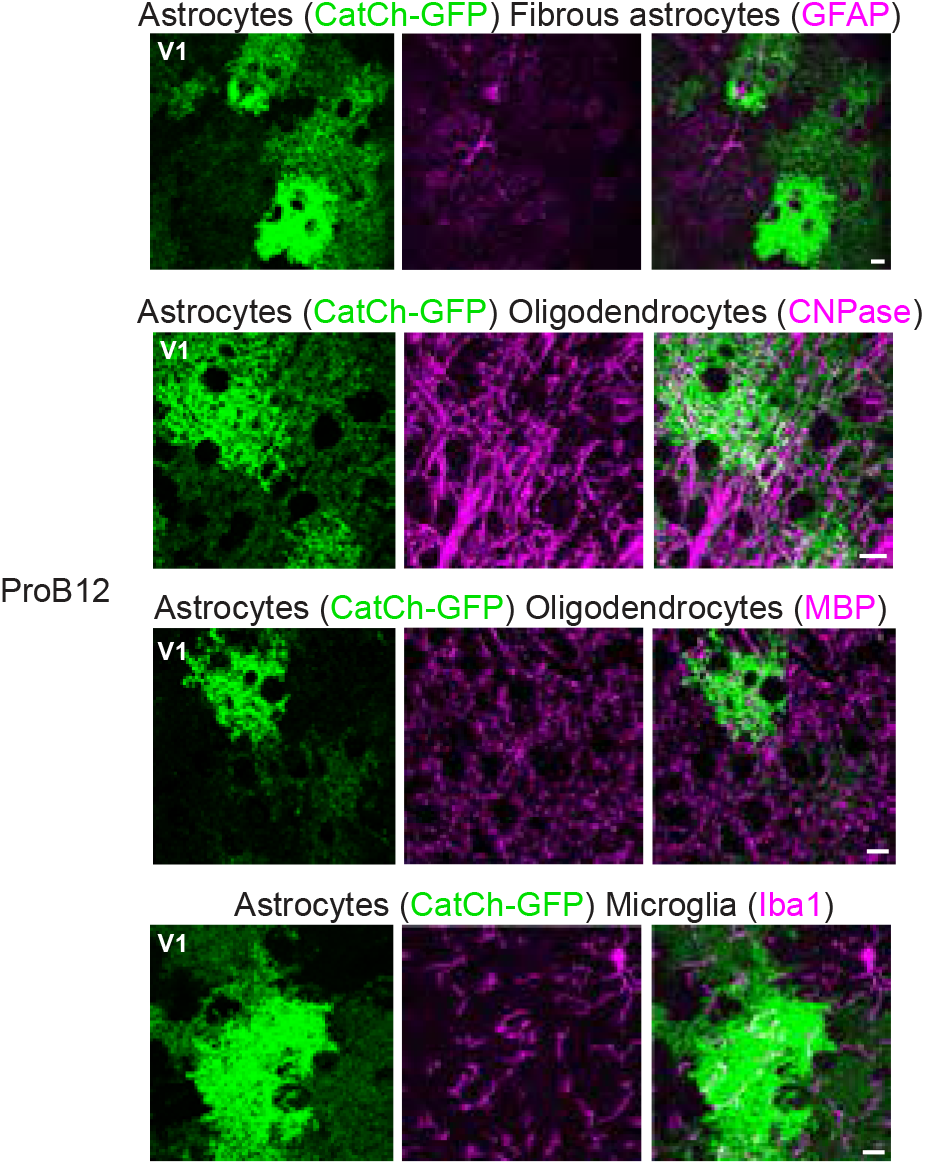
Astrocyte targeting by AAV-ProB12 in the visual cortex. Confocal images of AAV-infected V1. Left, CatCh-GFP (green); middle, immunostaining with marker (magenta) indicated above; right, CatCh-GFP and marker. Scale bars, 10 μm.

**Figure S4 related to Figure 5.**
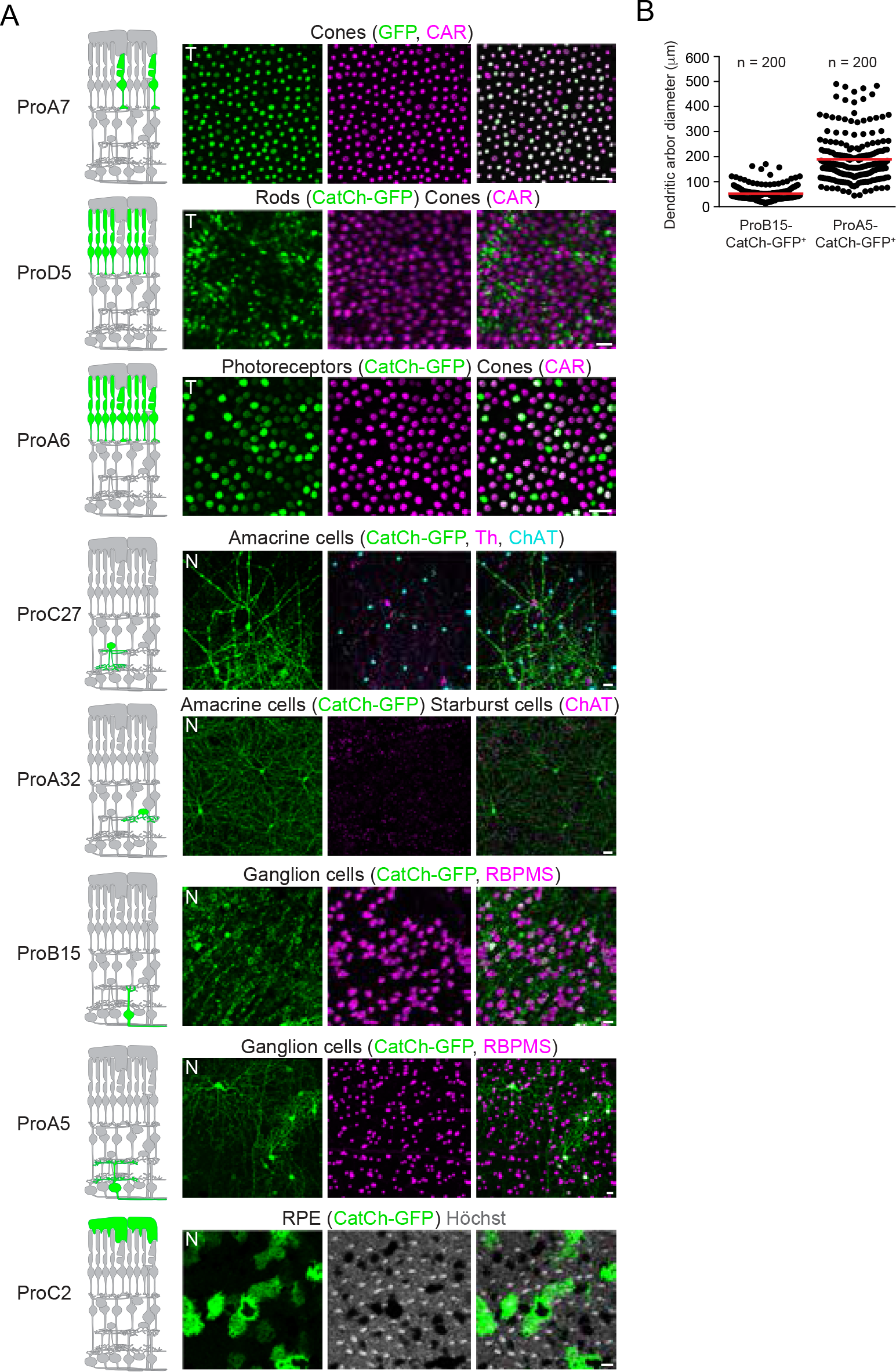
AAV-mediated cell-type targeting in non-human primate retina. (A) Confocal images of AAV-infected retinas (top view). Left, GFP or CatCh-GFP (green); middle, immunostaining with marker (magenta) indicated above or nuclear stain (Höchst, white); right, GFP or CatCh-GFP and marker or nuclear stain. Scale bars, 50 μm. (B) Quantification of the dendritic field diameter of cells targeted by AAV-ProB15 and AAV-ProA5 with means (red line) indicated.

**Figure S5 related to Figure 6.**
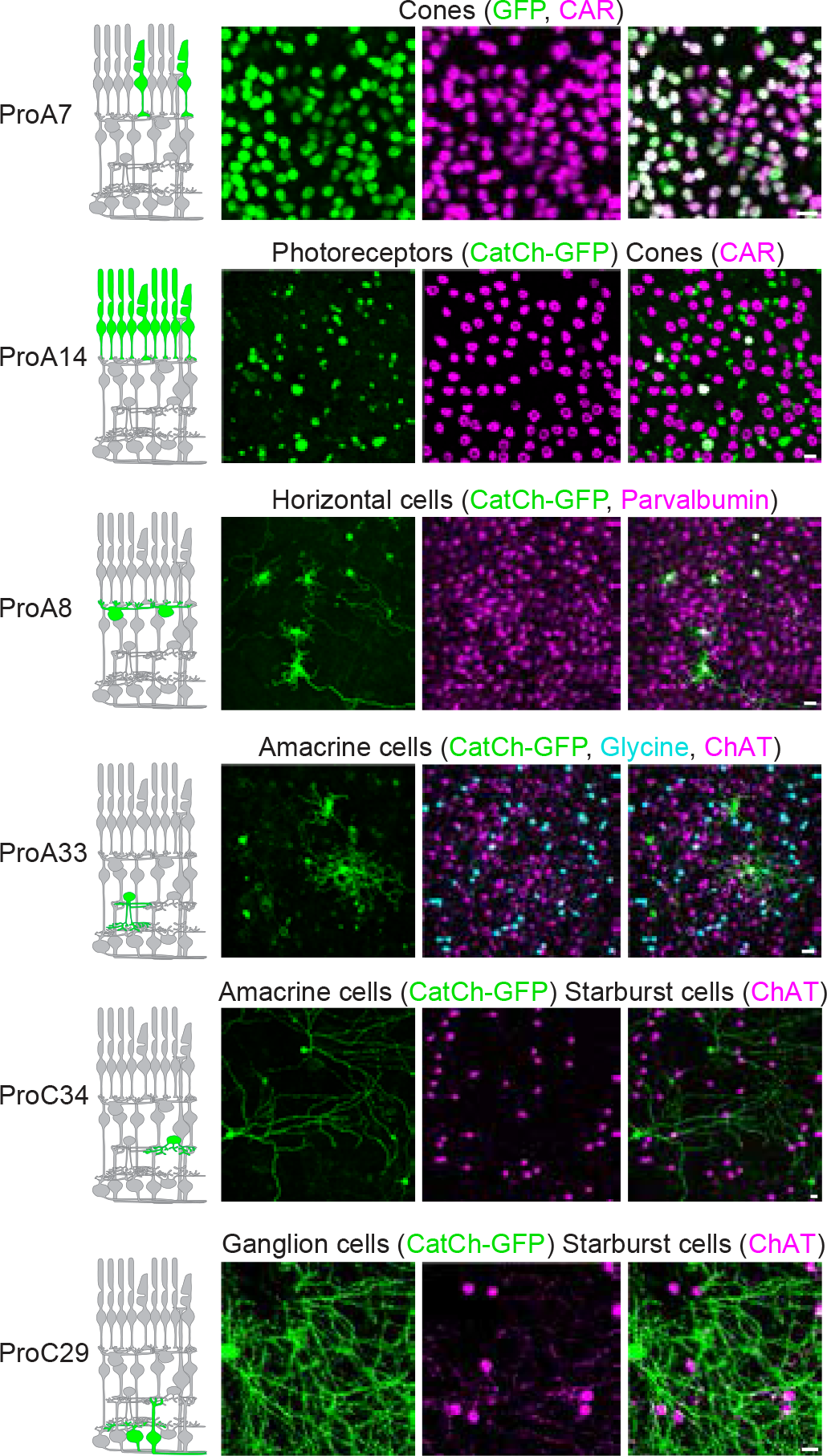
AAV-mediated cell-type targeting in human retina. Confocal images of AAV-infected retinas (top view). Left, GFP or CatCh-GFP (green); middle, immunostaining with marker (magenta) indicated above; right, GFP or CatCh-GFP and marker. Scale bars, 50 μm.

**Table S1.**
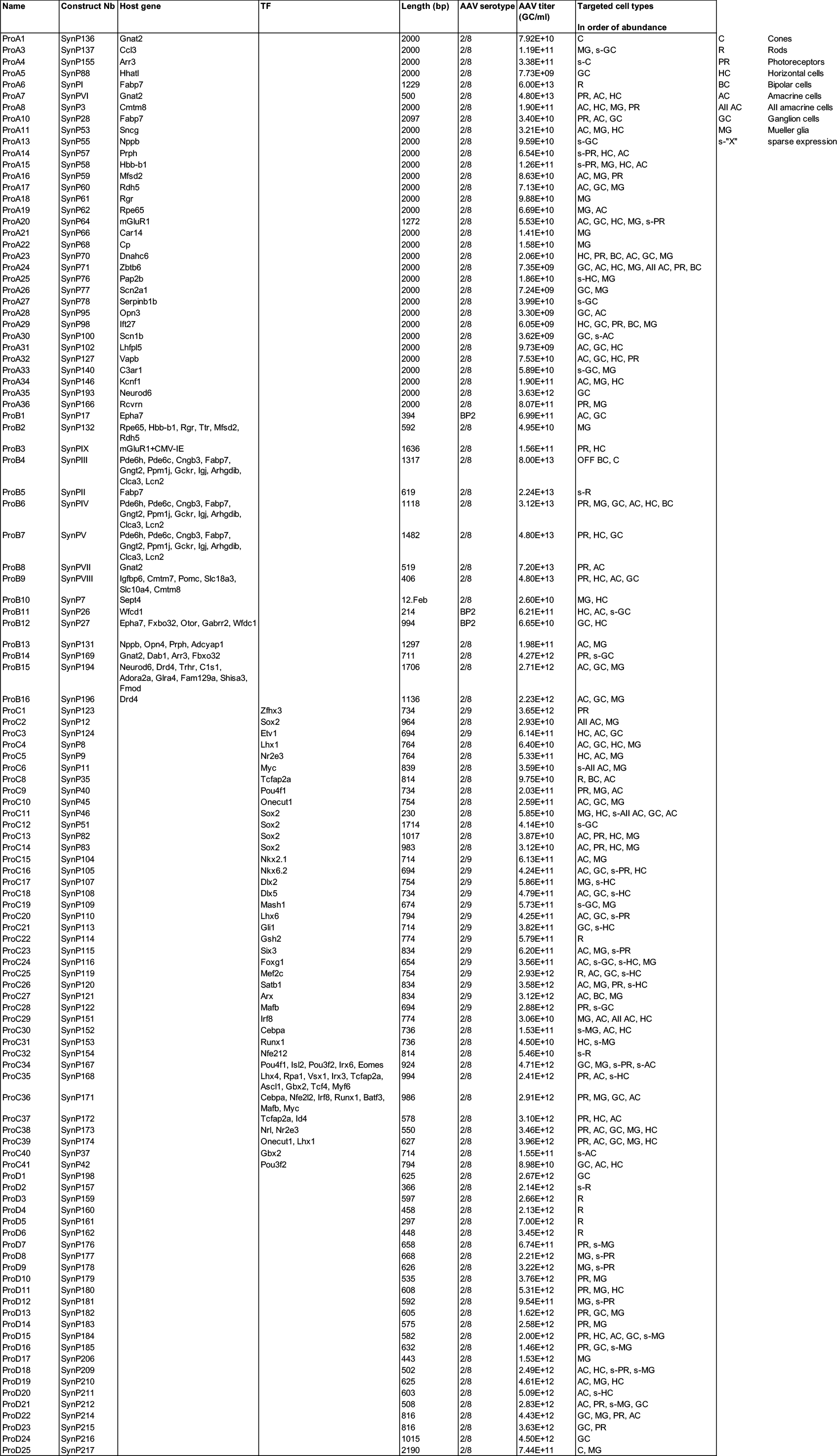
AAVs targeting mouse retinal cells.

**Table S2.**
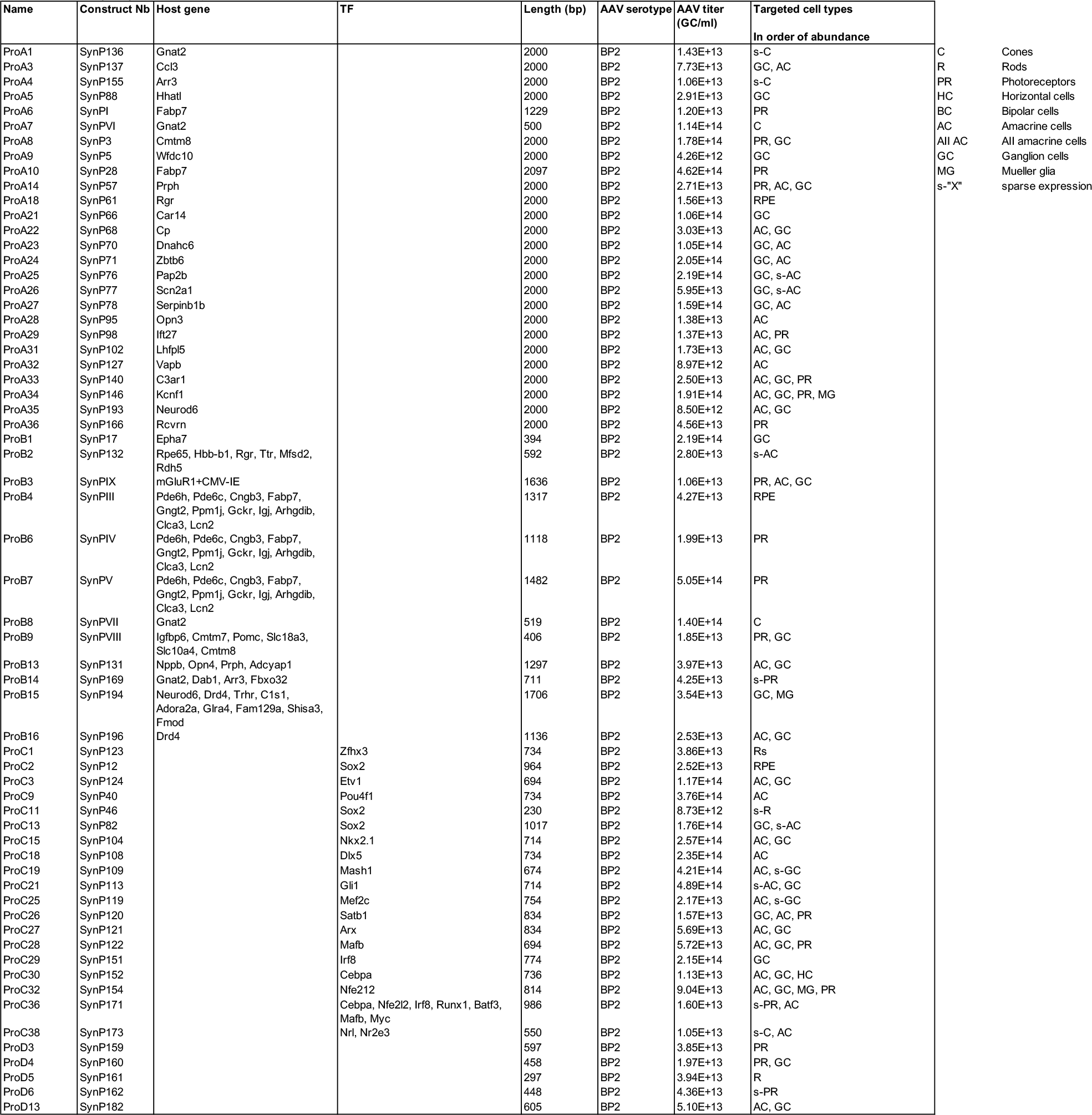
AAVs targeting non-human primate retinal cells.

**Table S3.**
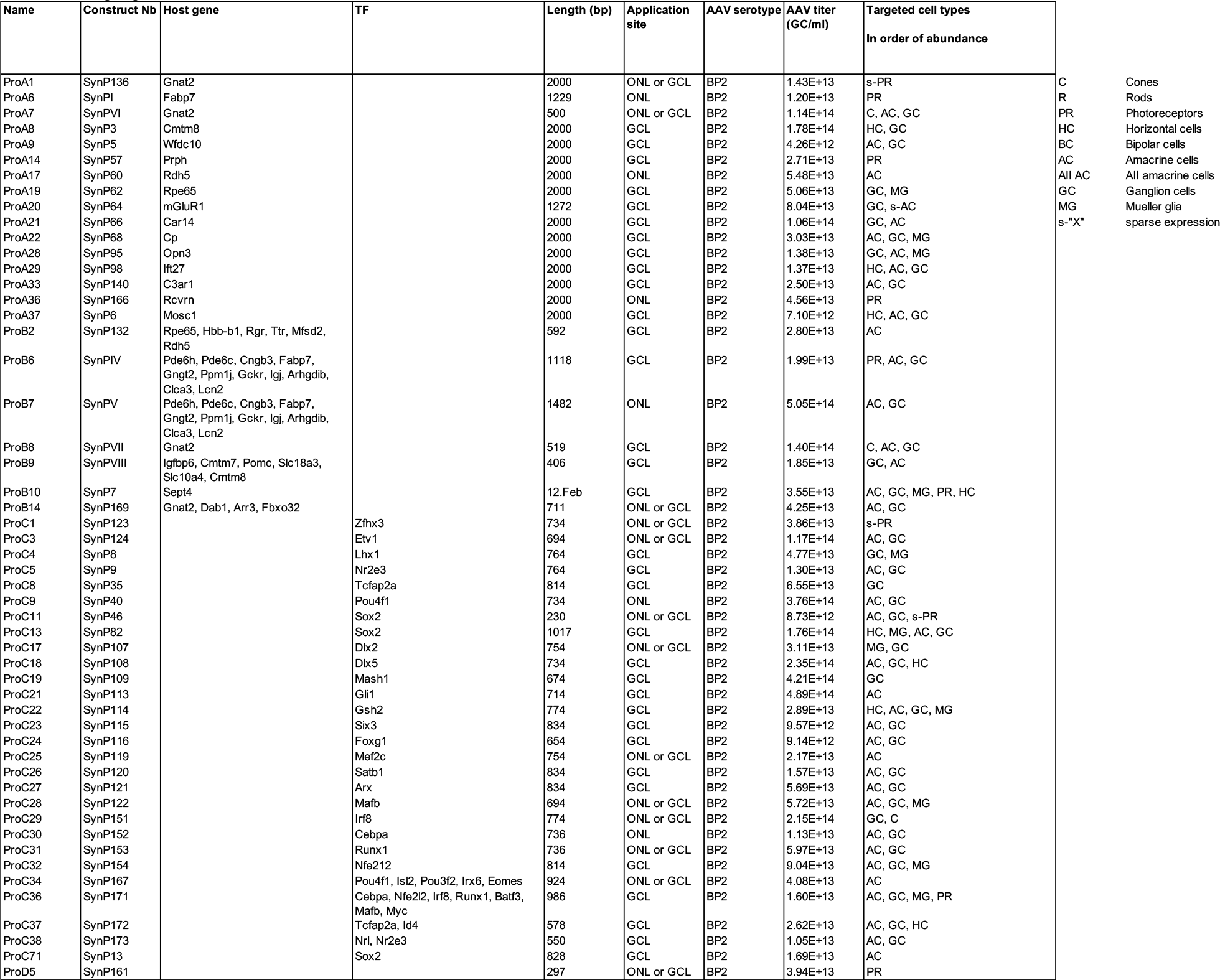
AAVs targeting human retinal cells.

